# Optogenetic Central Amygdala Stimulation is Highly Reinforcing and Strongly Outcompetes Fentanyl Self-Administration in Male, but not Female, Rats

**DOI:** 10.1101/2024.11.10.622812

**Authors:** E. Chaloux-Pinette, M. Vela Mirelez-Prince, S. Soto, K. Ananth, X. Tong, V. Silva, D. Rodriguez, P. H. Janak

## Abstract

The central nucleus of the amygdala plays a key role in many aspects of substance use disorders, in particular biasing behaviors toward one drug or natural reward over another, yet the role of this region in opioid self-administration remains unclear. Here, we report that pharmacological inactivation of the central amygdala reduces fentanyl self-administration, while intra-central amygdala opioid receptor antagonism dose-dependently increases fentanyl self-administration. We tested whether optogenetic activation of the central amygdala would increase motivation for fentanyl in rats with a long fentanyl self-administration history. While pairing fentanyl delivery with optogenetic central amygdala activation increased fentanyl self-administration, optoactivation itself was highly reinforcing and, in choice settings, was pursued over fentanyl despite mounting effort requirements, delays, and sporadic reward availability. Of note, under free response conditions, these effects were limited to male rats; while female rats avidly responded for optogenetic activation of the central amygdala, this activation was less effective in enhancing fentanyl intake, and it was not preferred over fentanyl. In contrast, under discrete trial choice which precluded independent regulation of intake of the two options, both females and males preferred optoactivation to fentanyl. These results demonstrate that optogenetic stimulation of neural activity within the anterior central amygdala does not appear to potentiate the reinforcing effects of fentanyl, but is itself highly reinforcing regardless of sex, and, in male rats can robustly outcompete fentanyl. In contrast, in females, fentanyl intake is relatively insensitive to competing opportunities for optoactivation, except when opportunities to obtain drug or optoactivation are sparse.

## Introduction

The central nucleus of the amygdala (CeA) is implicated in dependence, withdrawal, and relapse to drugs of abuse, with many findings indicating a prominent contribution later in the addiction cycle (Chaudun et al., 2024; de Guglielmo et al., 2016, 2019; Pelloux et al., 2018; Roura-Martínez et al., 2020; Venniro et al., 2020). Yet, evidence also implicates the CeA in acquisition and maintenance of drug intake (Berridge & Robinson, 2016; Fraser et al., 2024; Koob & Volkow, 2010). In rats, inactivation of the CeA decreases self-administration of cocaine, methamphetamine, and alcohol (Cain et al., 2008a; Cain et al., 2008b; X. Li et al., 2015a; W. L. Sun & Yuill, 2020a; Xue et al., 2012a; Warlow et al., 2017). Additionally, rats will self-administer amphetamine (Chevrette et al., 2002) and alcohol (Knight et al., 2020) directly into the CeA, implicating the region as a site for drug reinforcement. Similarly, studies have demonstrated involvement of the CeA in opioid self-administration(Cai et al., 2013a; Hou et al., 2017; Kallupi et al., 2020a), incubation of craving (Y. Q. Li et al., 2008), and withdrawal(Chaudun et al., 2024). Increased expression of GluA1 in the CeA increases morphine self-administration (Cai et al., 2013a; Hou et al., 2017) and intra-CeA infusion of the peptide nociception decreases oxycodone self-administration (Kallupi et al., 2020). In opioid dependent animals, mu-opioid receptor (MOR) expressing neurons within the CeA show enhanced activity during opioid withdrawal (Chaudun et al., 2024).

Optogenetic stimulation of neurons within the CeA is reported to bias choice behavior toward a drug or natural reward option (Robinson et al., 2014a; Tom et al., 2019a; Warlow et al., 2017, 2020a). Recently, when examining the impact of optogenetic activation during consumption of alcohol in a two-bottle choice procedure, we found that optogenetic activation of the CeA generally increases intake from the alcohol bottle paired with optostimulation as compared with a second alcohol bottle not paired with optostimulation (Fraser et al., 2024). Interestingly, some evidence suggests that intra-CeA activation of MOR might contribute to these types of motivational bias (Mahler & Berridge, 2009a). Collectively, these findings led us to examine contributions of the CeA to self-administration of the potent opioid, fentanyl, including in choice situations. We were particularly interested in studying both males and females, as the prior work discussed above focused on either females (Mahler & Berridge, 2009a; Robinson et al., 2014b; Tom et al., 2019b; Warlow et al., 2017, 2020b)or males (Fraser et al., 2024)), precluding direct comparison.

Here we report that CeA pharmacological inactivation in rats moderately decreases fentanyl self-administration, while naltrexone dose-dependently increases self-administration, supporting a role for this region in fentanyl’s reinforcing effects in both sexes. To further probe the role of the CeA in fentanyl self-administration, we explored whether CeA optostimulation can boost responding for fentanyl. We found optogenetic CeA stimulation increases fentanyl self-administration when paired with fentanyl infusions. Surprisingly, while CeA stimulation paired with fentanyl infusions biased choice over fentanyl alone in male and female rats, optostimulation itself was the preferred reinforcer in male but not female rats. To determine whether CeA stimulation increases motivation for fentanyl independent of the reinforcing properties of direct CeA stimulation alone, we assessed motivation for fentanyl and laser or simultaneous delivery of both reinforcers. In male rats, motivation for fentanyl paired with CeA stimulation, while greater than for fentanyl alone, was not significantly greater than motivation for CeA stimulation alone.

Female rats did not display significantly different levels of motivation for fentanyl, fentanyl paired with CeA stimulation, or CeA stimulation. Male and female rats persisted in pursuing CeA stimulation despite increasing effort requirements, delays in delivery of stimulation, and limited reward availability. These results demonstrate CeA stimulation is highly reinforcing and is preferred over fentanyl in male rats with extensive fentanyl self-administration training.

## Methods

### Subjects

Adult male and female Sprague Dawley rats (ENVIGO, Frederick, MD) weighing 250-300g and 200-225g, respectively, at the time of surgery were used in all experiments. Rats were single-housed in a temperature and humidity-controlled vivarium on a normal light cycle (12/12h light/dark cycle, lights on at 7am) with *ad libitum* access to food and water. Rats were trained during the light cycle. The procedures used were approved by the animal care and use committee of Johns Hopkins University and were conducted in accordance with the National Institutes of Health Guidelines for the Care and Use of Laboratory Animals.

### Drugs

Fentanyl Citrate (Sigma-Aldrich or Cayman Chemical) was diluted in 0.9% sterile saline at 10µg/ml. Muscimol hydrobromide and Baclofen hydrochloride (Sigma-Aldrich) were diluted in 0.9% sterile saline to a final concentration of 195mg/ml Muscimol and 2130mg/ml Baclofen (0.1mM/1mM). Naltrexone hydrochloride (Sigma-Aldrich) was diluted in 0.9% sterile saline to concentrations of 1.33, 4.17, 13.33, or 41.67 mg/ml. Brevital sodium (Henry Schein) was diluted to 10mg/ml with 0.9% sterile saline.

### Surgeries

#### Catheter implantation

Rats were anesthetized with isoflurane (3-5% induction, 1-2% maintenance) and administered the analgesic rimadyl (5mg/kg, SC, Zoetis) and the antibiotic cefazolin (70mg/kg, SC, West-Ward) prior to catheter implantation. For microinfusion experiments, silastic jugular catheters were constructed as described previously (Thomsen & Caine, 2005). For optogenetic experiments, animals were implanted with a single channel vascular access button connected to a polyurethane catheter (Instech Labs; Plymouth Meeting, PA USA). The catheter tubing was threaded under the skin from a subcutaneous base in the mid-scapular region and inserted into the right jugular veinClick or tap here to enter text.. Catheters were flushed daily with ∼0.1ml solution of 10mg/ml gentamicin sulfate (VetOne) and 100 IU/ml Heparin (Sagent) in sterile saline. Once behavioral training began, catheters were flushed before and after self-administration and daily on rest days and were assessed daily for blood return. If blood return was absent, rats were administered 0.1ml of brevital sodium (10mg/ml in sterile saline, Henry Schein). If the rat did not become ataxic within 10 seconds, the catheter was considered not patent and the rat was removed from the study or a new catheter was implanted into the left jugular vein.

#### CeA cannulation

Immediately after catheterization, anesthetized rats were implanted with bilateral cannula targeted to the CeA using standard stereotaxic procedures. A 22-gauge guide cannula (P1 technologies; Roanoke, VA) was implanted above the CeA in each hemisphere (for males: AP: -2.4 mm, ML ± 4.2 mm, DV −7.0 mm; for females: AP: -2.4 mm, ML ± 4.2 mm, DV −7.0 mm; all coordinates relative to bregma). In one cohort of animals, the cannulas were implanted with DV: -6mm due to use of a 22-gauge cannula with shorter projection. To prevent cannula occlusion, stainless steel stylets were placed in the cannulas after surgery and only removed during micro-infusions. Rats were allowed to recover for at least a week prior to behavioral training.

#### Optogenetics

Immediately after catheterization, anesthetized rats received virus infusion and optical fiber placement using standard stereotaxic procedures. Rats received bilateral infusions of AAV ChR2 (AAV5-Hsyn-ChR2-eYFP; Addgene; Watertown, MA) in the CeA (AP: −2.4, ML: ±3.9-4, DV: −7.8; N=6 F, 7M). A total of 0.5µl of virus was infused per side over 5 minutes at a constant rate (0.1µl/min). The injector was left in place for an additional 10 minutes to allow for diffusion. Additional rats were infused with AAV eYFP (AAV5-Hsyn-eYFP; Addgene; Watertown, MA) to serve as inactive virus controls. After viral infusion, rats were implanted with bilateral optic fibers (300µm) aimed 0.3mm dorsal to the infusion sites. Rats were allowed to recover for at least a week prior to behavioral training. Optogenetic manipulations began a minimum of 4 weeks after viral infusion.

### Self-administration training and microinfusions

#### Self-administration training

Rats were trained in standard Med Associates operant chambers housed inside sound-attenuating chambers (Med Associates, St Albans, VT, USA). Operant chambers were fitted with two retractable levers (active and inactive), stimulus lights above each lever, speakers, and a house light. Boxes were controlled using Med-PC IV software (Med Associates, St Albans, VT, USA).

Rats were trained to press one lever (active lever) on a fixed-ratio 1 (FR1) schedule in 2-hour sessions ∼6 days per week. An inactive lever was available for the naltrexone dose response and loading dose experiments and was included for a subset (n=7 female, 6 male) of the rats micro-infused with saline, M/B and naltrexone. The stimulus light above the active lever indicated drug availability. Active lever presses resulted in delivery of 0.5µg fentanyl in 50µl sterile saline delivered over 2.8 seconds (Fragale et al., 2020). Drug delivery was accompanied by a 2.8s tone and was followed by extinguishing of the stimulus light and illumination of the house light for a 20 second timeout. After 20 seconds elapsed, the house light extinguished, and the stimulus light was re-illuminated to indicate availability of drug. Training occurred for at least 28 sessions after which the contribution of CeA activity or CeA opioid receptors to fentanyl self-administration was assessed. Training data and the locations of the infuser tips for rats included in this study are shown in Figure 1.

**Figure 1.**
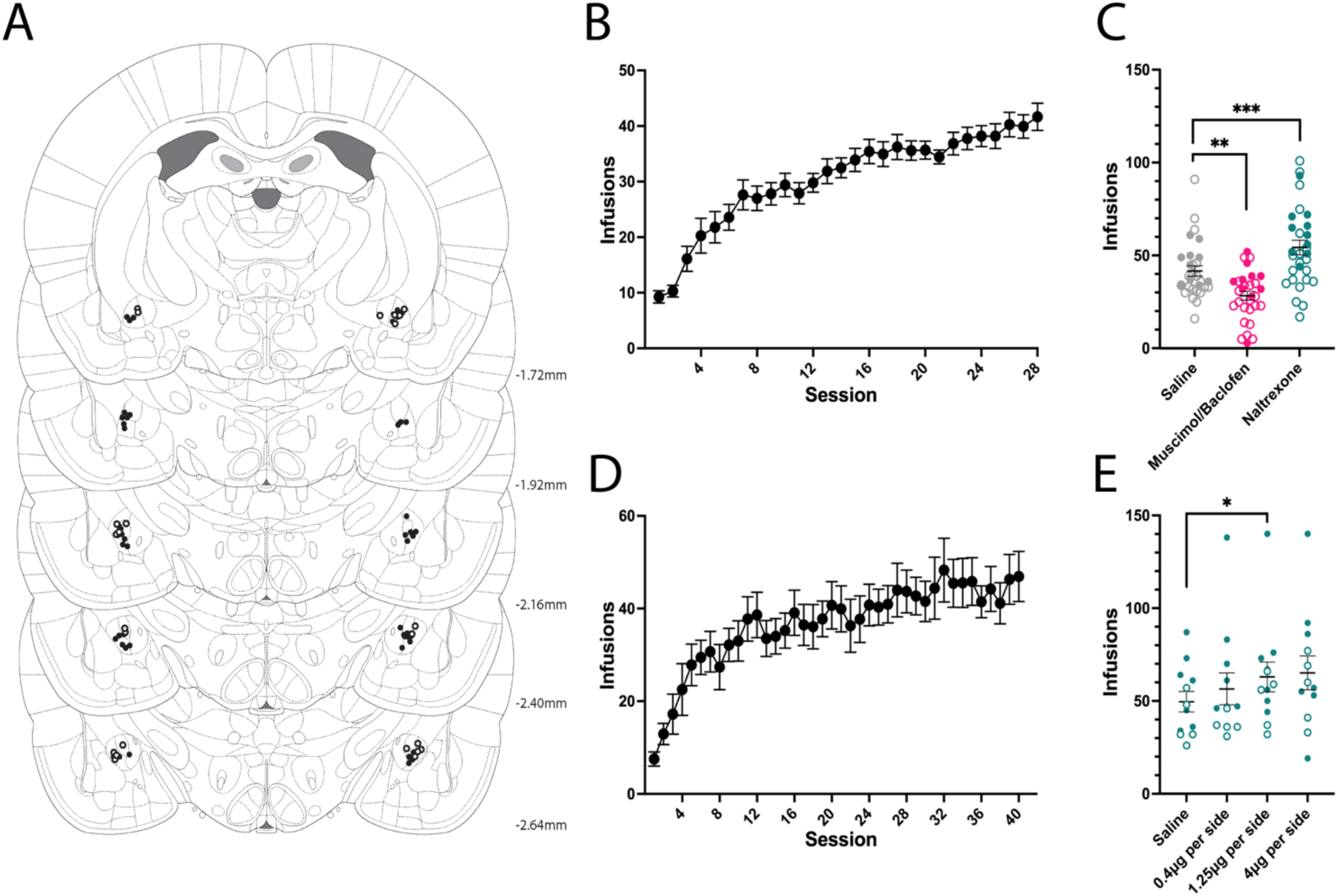
CeA pharmacological inactivation and opioid receptor antagonism differentially effect fentanyl self-administration. (A) Black circles represent approximate locations of infuser tips for(•) cohort 1 and (o) cohort 2 (B) Cohort 1 training session data n=30 (20 male, 10 female). (C) Effect of CeA pharmacological inactivation and opioid receptor antagonism on total fentanyl self-administration (D) Cohort 2 training session data n=12 (5 male, 7 female). (E) Effect of naltrexone infused into the at different doses on total fentanyl self-administration Data are presented as mean± SEM. * indicates significant main effect of treatment or significant pairwise difference, * p<0.05, ** p≤0.01, *** p≤0.001, **** p≤0.0001.

#### Microinfusions

Rats were assigned to balanced groups for CeA micro-infusion, based upon average fentanyl infusions over the last 3 training days. Rats received naltrexone (0.4 µg, 1.25µg, or 4 µg/hemisphere, Sigma-Aldrich, St. Louis, MO), a mixture of the GABA_A_ receptor agonist, muscimol, and GABA_B_ receptor agonist, baclofen, (5.85ng and 64.10ng per hemisphere, respectively; Sigma-Aldrich, St Louis, MO), or saline vehicle prior to self-administration sessions. Infusions into the CeA were made in a volume of 0.3µl over 60 seconds via 28-gauge infusers (P1 technologies; Roanoke, VA) connected via PE50 tubing to 10 µl Hamilton syringes (Hamilton Company; Reno, NV) secured in a motorized pump (Harvard Instruments; Holliston, MA). Infusers extended 1 or 2mm below the guide cannula (final DV: -8mm) depending on guide cannula length. During infusions, rats were held gently and infusion volume was monitored to ensure that all rats received the full infusion volume. Infusers were removed and stylets replaced one minute following the infusion. Rats were tested for fentanyl self-administration 10 minutes following drug infusion.

### Training and behavioral procedures for optogenetic experiments

#### Fentanyl self-administration training

Rats were trained in standard Med Associates operant chambers housed inside sound-attenuating chambers (Med Associates, St Albans, VT, USA). Operant chambers were fitted with three retractable levers (two active levers, equally spaced along one wall, and one inactive lever placed on the opposite wall), stimulus lights above each active lever, speakers, and a house light. Boxes were controlled using Med-PC IV software (Med Associates, St Albans, VT, USA).

Rats were trained to lever-press on a fixed-ratio 1 (FR1) schedule for 4-9 sessions, FR2 for 1-2 sessions, and then FR3 until behavior showed less than 20% variability between sessions (average 41.38 total sessions, range: 40-42). The stimulus light above the active lever indicated drug availability. Ratios completed on the active lever resulted in delivery of 0.5µg fentanyl dissolved in 50µl sterile saline delivered over 2.8 seconds. Drug delivery was accompanied by a 2.8s tone, after which the stimulus light was extinguished, and the house light was illuminated for a 20 second timeout. After 20 seconds elapsed, the house light extinguished, and the stimulus light was re-illuminated to indicate availability of drug. Both the active and inactive levers were available for the duration of the session. After training rats progressed to optogenetic tests described below. All rats had at least three sessions to acclimate to being tethered via both patch cord and catheter tubing before any optogenetic testing occurred. Unless otherwise stated, all sessions were 2 hours long.

#### Fentanyl infusion-paired CeA optostimulation

Conditions were identical to FR3 self-administration training with the exception that FR3 completion resulted in both infusion of 0.5µg fentanyl and an 8s blue (473nm) light stimulation (7-10mW 25Hz) of the CeA bilaterally (8s laser).

#### Non-contingent inter-infusion interval CeA optostimulation

Rats self-administered fentanyl under an FR3 response requirement. For each subject we determined an average value for the 25th percentile and 75th percentile inter-infusion intervals according to 5 fentanyl self-administration sessions prior to the beginning of testing and a value within that range was randomly selected at the end of each timeout. If that interval passed without a lever press, rats received an 8 second blue (473nm) light optostimulation (25HZ, 7-10mW). If a lever press occurred, the timer reset for the interval (i.e. the entire interval was required to pass between the LP and the stimulation). If a reward was earned before the interval elapsed, no optostimulation occurred and a new interval was selected at random at the end of the timeout period. Consequently, stimulation did not occur during all inter-infusion intervals. For comparison, the baseline session was run through a custom MATLAB script that determined “stimulation” times in an identical manner. This program operated on the same logic as the test, producing a collection of “stimulation” times and response data for those periods.

#### Choice between two rewards

Rats underwent choice sessions where two distinct rewards were available out of three possible rewards: fentanyl, 0.5s laser, or fentanyl accompanied by 0.5s laser. Each choice session began with two forced-choice trials during which active levers were presented one at a time. This allowed rats to sample the possible outcomes. The inactive lever was extended for the entire session. Following the forced choice trials, both active levers extended and the lights above them were illuminated to indicate reward availability. Once an FR3 was completed for either active lever, a tone sounded for 2.8 seconds. Tones (2,900 and 4,500 Hz) were assigned to each lever and were independent of reward identity during all testing. Lights above the levers were extinguished during the timeout period and a house light was illuminated. Responses on either active lever during the 2.8 second tone or the 20 second timeout had no effect. There was at least one fentanyl self-administration session between choice tests and both test order and lever assignments were counterbalanced across rats.

#### Progressive ratio responding for fentanyl and optostimulation rewards

Maximum effort and motivation to seek each reward was measured by response breakpoint. Each session had only one reward available (fentanyl, 0.5s laser or fentanyl + 0.5s laser). Responses required on active lever increased exponentially after each reward (1, 2, 4, 6, 9, 12, 15, 20, 25, 32, 40, 50, 62, 77, 95, 118, 145, 178, 219, 268,…) according to the formula PR = [5e(reward number × 0.2)] – 5 and rounded to the nearest integer2–4. The session continued until 30 minutes had passed without active lever responses or after 4 hours had elapsed.

#### Choice between CeA optostimulation and drug reward when response requirement increases for one option

Rats were presented with either fentanyl vs. 0.5s laser or fentanyl+0.5s laser vs. 0.5s laser. One of the two rewards was under a PR response requirement on one lever while the other remained available under an FR3 on the other lever. Order of choice pairings and response requirement was counterbalanced across animals. Rats had at least one fentanyl self-administration session between the two sessions with the first reward pairing and the two sessions for the second reward pairing. Each choice session began with two forced-choice cycles (four total trials) with FR3 response requirements. After the forced choice trials, the PR lever required an increasing number of lever presses according to the distribution described above. Response requirement reset once a reward was earned on the FR3 lever and lever presses during the reward delivery and timeout period (22.8s) were not counted towards the response requirement. During this 22.8 second period, rewards could not be earned on either lever. The session continued until 30 minutes of no activity on the PR lever or after 4 hours had elapsed.

#### Persistence in choosing increasingly delayed CeA optostimulation versus fentanyl

To assess the relative value of the 0.5s laser reward versus fentanyl, rats underwent a delay breakpoint task where delivery of the laser reward was delayed progressively longer. All sessions began with two forced-choice cycles as above, with no delay in delivery of laser reward. Following these trials, all levers remained extended and delivery of the laser was delayed by 0s after the first laser reward and increased by 1s for every subsequent laser reward. Reward delivery was accompanied by a 2.8s tone followed by a 20s timeout during which lights above active levers were extinguished and a house light was illuminated. Delays experienced were dependent on the number of laser rewards earned.

#### Discrete choice behavior with extended ITI

We next wanted to assess the relative value of 0.5s laser reward versus fentanyl when total rewards and reward frequency were constrained. In place of the 20s timeout, a 3min intertrial interval (ITI) was imposed. Each session began with two “forced choice” cycles with a 3min ITI, after which both levers were extended. Once the response requirement was completed on either lever, the reward was delivered and a 2.8s tone sounded. Both active levers retracted after the tone. For the 3min ITI, lever lights were extinguished, and the house light was illuminated. Rats were given 20 free choice trials. If they did not earn a reward in 5min during free choice sessions, the trial was considered an omission, the levers retracted and the 3min ITI began.

#### Intracranial self-stimulation

During intracranial self-stimulation sessions two nose-poke ports were available. The house light was illuminated for the duration of the session. A nose-poke in the active port resulted in the delivery of 0.5s laser stimulation at either 0.5-1mW or 7-10mW power. The active and inactive ports were alternated across sessions. Nose-pokes and reward delivery were not accompanied by any cues, nor was there any cue distinguishing active from inactive port. Sessions lasted 45 minutes and optostimulation was delivered either unilaterally or bilaterally.

Fentanyl-naïve animals underwent additional self-stimulation sessions with FR3 lever press response requirements to earn 500ms or 8s stimulation at 7-10mW power.

### Histology

#### Assessment of cannula placement

Following completion of the experiment, brains were extracted and post-fixed in 4% paraformaldehyde in 0.1M NaPB for 48 hours and then cryoprotected in 30% sucrose in 0.1M NaPB. Brains were sectioned coronally (40 µm) on a cryostat, mounted, and stained with Cresyl violet (FD Neurotechnologies; Ellicott City, MD). Micro-infusion sites were mapped onto a rat brain atlas (Paxinos & Watson, 2009).

#### Assessment of viral expression and optic fiber placement

Rats were anesthetized 90 minutes after the final ICSS session with an overdose of sodium pentobarbital (70-150mg, IV) and transcardially perfused with cold 1M PB followed by cold 4% PFA. Brains were extracted and stored in 4% PFA for 24 hours before being transferred to a solution of 30% sucrose in 1M PB. Brains were coronally sectioned (40µm) on a cryostat, sections were then mounted on slides and cover slipped with Vectashield HardSet mounting medium with DAPI (Vector Laboratories; Newark, CA USA). Slides were then imaged using a Leica fluorescent microscope (Figure 2). Results were marked using the rat brain atlas (Paxinos & Watson, 2009).

**Figure 2.**
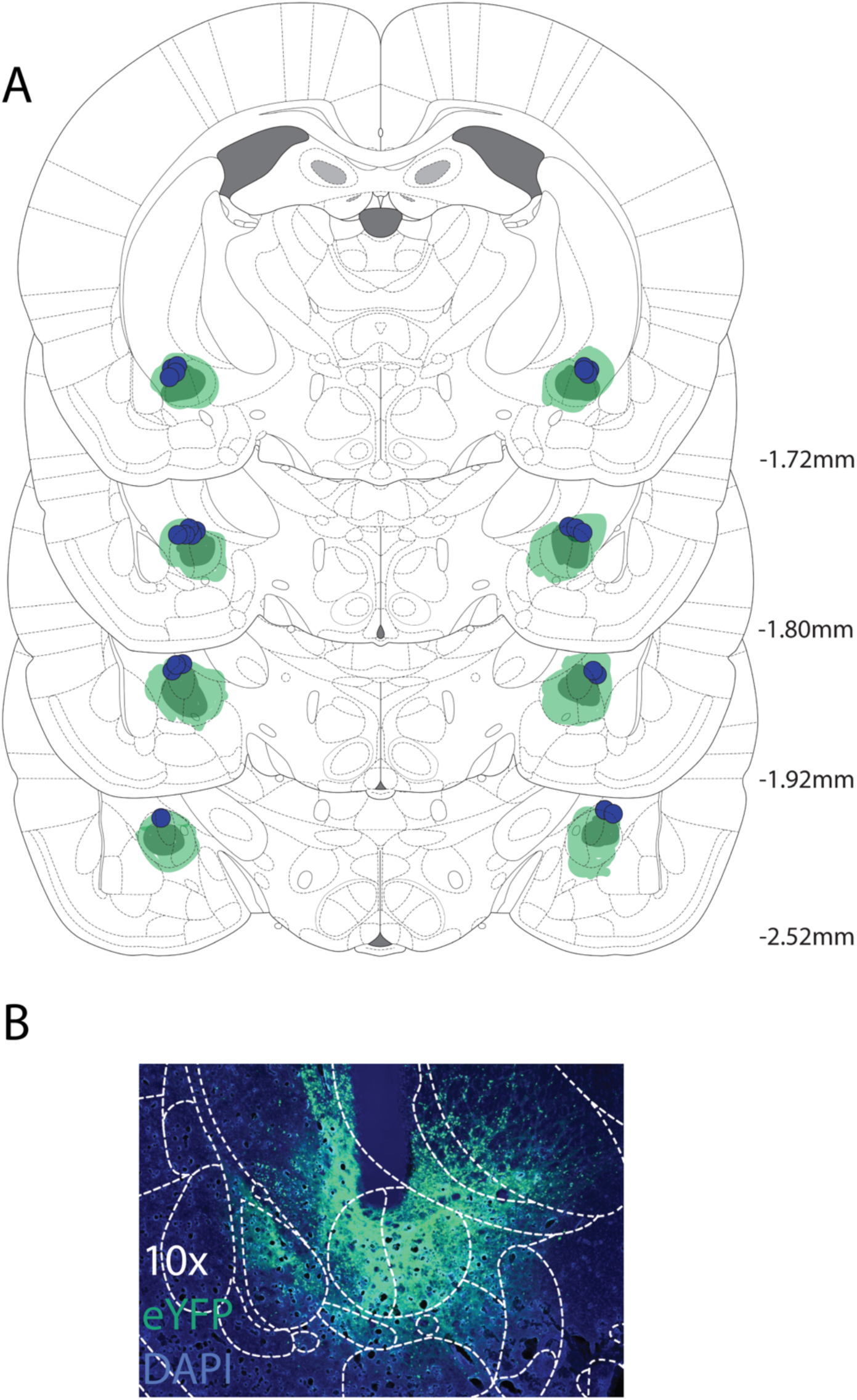
Viral expression and optic fiber placements in rats undergoing optogenetic stimulation of the CeA (A) Optic fiber tip locations are denoted by blue circles, in addition to approximate maximal (dark green) and overall (light green) viral expression. (B) Example images of expression of ChR2-eYFP within the central amygdala (green) and nuclear staining with DAPI in blue.

### Statistical analysis

Data were analyzed with GraphPad Prism 10 (La Jolla, CA). A significance level of α=0.05 was used for all analysis. Geisser-Greenhouse correction was applied in ANOVAs where sphericity could not be assumed. For microinfusion experiments there was no significant main effect of sex or interaction between sex and test conditions. Consequently, for those experiments data were collapsed into a mixed-sex data set for all further analysis. When the ANOVA revealed an interaction, pairwise comparisons were performed using Dunnett’s multiple comparison test. Data from optogenetics experiments were analyzed using two- or three-way/repeated measures ANOVAs or Pearson’s correlation, as appropriate. For optogenetics experiments, sex was considered as a between subjects factor. Reward choice and/or session were considered as within-subjects factors. All pairwise comparisons were done using Sidak’s multiple comparison test. When data were non-normal, non-parametric analyses were used, and correlation analyses were done by computing Spearman correlation coefficient.

## Results

### CeA inactivation decreases and opioid receptor blockade increases fentanyl self-administration

Here we tested the effect of temporary inactivation and of opioid receptor blockade on fentanyl self-administration. Following fentanyl self-administration training (Figure 1B), rats received CeA micro-infusions of muscimol and baclofen (MB), naltrexone, or saline in a counterbalanced order 10 minutes prior to the start of a self-administration session. A 2-way ANOVA revealed a significant main effect of treatment on fentanyl intake, as measured by number of infusions earned (Figure 1C, F_(1.23,34.43)_=29.57, p<0.0001), accounted for by a decrease fentanyl self-administration after reversible pharmacological inactivation of the CeA (Figure 1C, p=0.001), and an increase after naltrexone (1.25µg/hemisphere; Figure 1C, p<0.0001). There was no significant effect of sex (F_(1,28)_=3.457, p=0.0735) or interaction between sex and treatment effects (F_(2,56)_=29.57, p=0.557).

To further explore the effect of CeA naltrexone on responding for fentanyl, an additional cohort of rats (Figure 1D, n=12, 5 male, 7 female) was trained to self-administer fentanyl and then was administered various doses of naltrexone (0, 0.4, 1.25, or 4µg/hemisphere) into the CeA prior to self-administration sessions, again revealing a moderate effect on number of infusions (Figure 1E; Friedman statistic=11.72, p=0.008). While 0.4 µg failed to significantly alter fentanyl intake (p>0.999), and the effect of 4µg of naltrexone did not reach significance (p=0.0632), there was a significant increase in the number of infusions after 1.25µg naltrexone (p=0.049).

### CeA optogenetic stimulation during fentanyl delivery increases self-administration in males, but not females

Our findings above indicate that the CeA neural activity can regulate fentanyl self-administration. Prior studies found that optogenetic CeA activation enhances responding for natural reward and cocaine (Robinson et al., 2014b; Warlow et al., 2017, 2020b) and enhances consumption of alcohol and natural reward (Fraser et al., 2024). Therefore, we assessed the impact of reward-paired optogenetic activation of CeA neuronal activity in rats trained to self-administer fentanyl on an FR3 schedule. Prior to initiation of optogenetic manipulations, we examined the training data for sex differences. We found no sex difference in number of infusions late in acquisition after responding had plateaued (Figure 3A, Sex: F_1,11_=0.380, p=0.550; Sex x Session: F_19,209_=0.548, p=0.937), although male rats were on average more variable in their responding (coefficient of variation analysis; Figure 3B, Mann-Whitney test: p=0.035).

**Figure 3.**
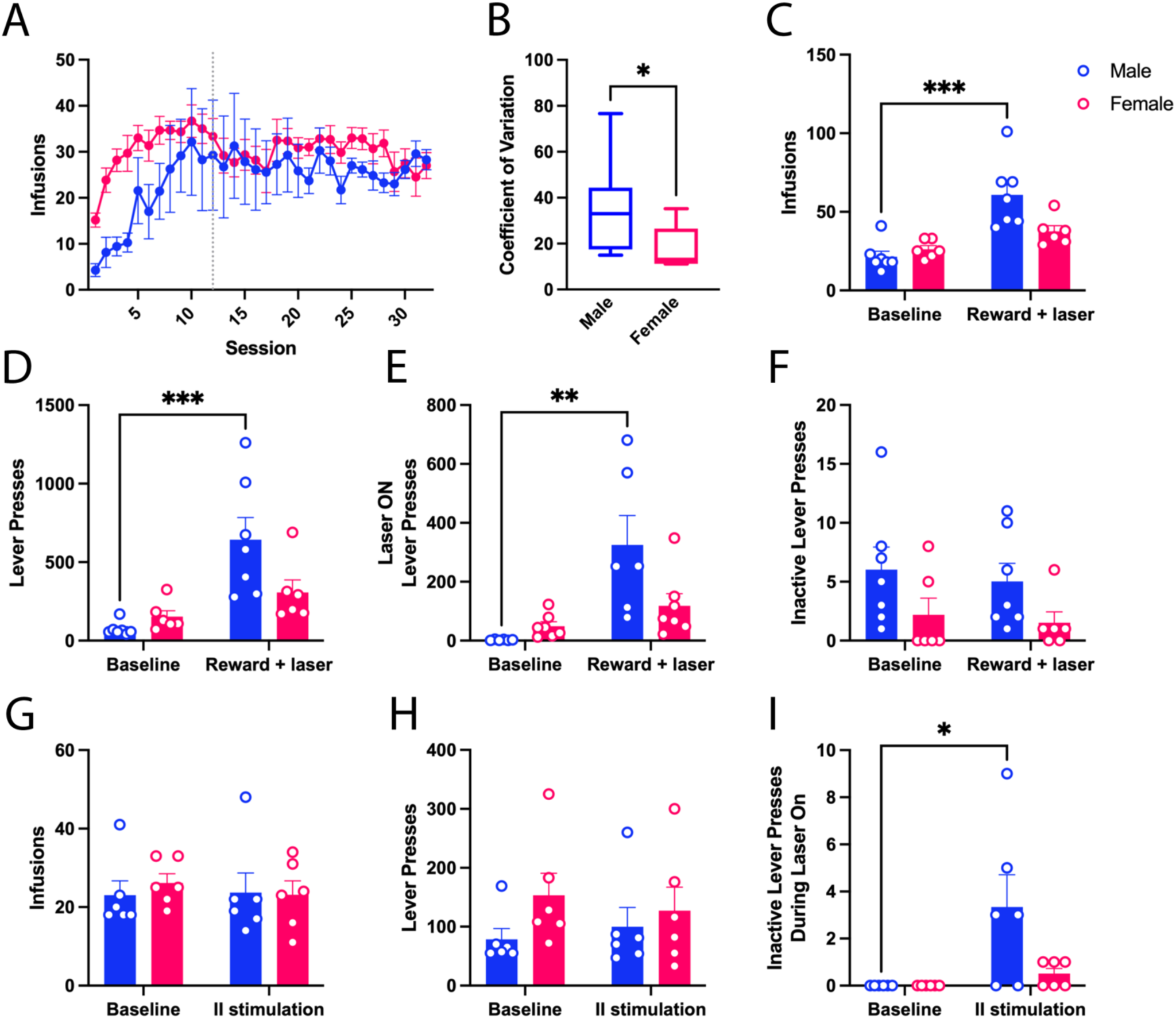
Optogenetic stimulation of the CeA during fentanyl infusion increases fentanyl self-administration in males. (A) Training data for females and males. Dotted line denotes session 12. (B) Coefficient of variation for fentanyl intake during fentanyl self-administration sessions post-acquisition. (C-F) comparison of responses between baseline session and session pairing CeA stimulation and fentanyl infusion. (C) infusions, (D) lever presses, (E) lever presses during stimulation, and (F) inactive lever presses. (G-1) comparison of responses between baseline session and session with inter-infusion interval CeA stimulation (II Stimulation) (G) infusions, (H) total active lever presses, and (!)inactive lever presses during stimulation period. Data are presented as mean ± SEM. N=6 female, 7 male *p<0.05, **p≤0.01, ***p≤0.001.

Next, rats were allowed to self-administer fentanyl with completion of the response requirement paired with delivery of an 8 second blue (473nm) light optogenetic stimulation (Click or tap here to enter text.25HZ, 7-10mW) simultaneous with the delivery of the 3-second tone stimulus and fentanyl infusion. Optostimulation at the time of reward delivery increased the number of fentanyl infusions received per session (3C, main effect of Stimulation: F_1,11_=19.83, p=0.001), but selectively in males (Sex x stimulation interaction: F_1,11_=6.076, p=0.031; baseline vs stimulation session. males: p=0.0007, females: p=0.364). Analysis of the number of lever press responses made to obtain fentanyl revealed a similar pattern; optostimulation increased active lever pressing both throughout the session (**Error! Reference source not found.**D, main effect of Stimulation: F_1,11_=16.07, p=0.0023) and during the 8-second laser stimulation period (Figure 3E, main effect of Stimulation: F_1,11_=14.33, p=0.003), with significant increases again only in males (Sex x stimulation interaction: Total active lever presses: F_1,11_=16.07, p=0.0413; Lever presses during stimulation: F_1,11_=6.015, p=0.0323). Specifically, males showed increased active lever pressing overall (Figure 3D, p=0.0014) and during laser stimulation (Figure 3E, p=0.003), while active lever pressing was not significantly affected in females (Overall: p=0.570; Stimulation period: p=0.574). There was no effect of stimulation on inactive lever pressing (Figure 3F, Stimulation: F_1,11_=0.3807, p=0.550), suggesting that the change in behavior with reward-paired CeA stimulation was not due to nonspecific motor activation.

To test whether the increase in self-administration by CeA stimulation depends on stimulation at the time of the response and drug infusion, we examined the impact of non-contingent CeA stimulation to influence fentanyl self-administration behavior, by delivering CeA optostimulation during the inter-infusion interval. Non-contingent CeA stimulation did not alter fentanyl self-administration or drug lever pressing during the session (Figure 3G, main effect of Stimulation: Infusions: F_1,10_=0.700, p=0.467; Drug lever presses: F_1,10_=0.085, p=0.777), and there were no differences on infusion numbers relative to sex (Sex: F_1,10_=0.067, p=0.801; Sex x Stimulation: F_1,10_=1.729, p=0.218), however there was an interaction between sex and stimulation on number of drug lever presses (Figure 3H, Sex: F_1,10_=1.226, p=0.294; Sex x Stimulation: F_1,10_=7.274 p=0.0224), although none of the individual comparisons were statistically significant. Similarly, we found no main effect of stimulation on active lever presses and infusions delivered during the 8-s optostimulation period itself relative to no-stimulation baseline (Data not shown, main effect of Stimulation: Infusions: F_1,10_=0, p>0.999; Drug lever presses: F_1,10_=0.01259, p=0.913). However, rats made more inactive lever presses during the inter-infusion stimulation period than the baseline “stimulation” periods (Figure 3I, Stimulation: F_1,10_=5.859, p=0.0209). No effect of laser was observed in an eYFP control group (n=5) that did not express ChR2 (supplemental figure S1A-B).

### CeA optostimulation is highly reinforcing and outcompetes fentanyl as a reinforcer

The results above show that CeA optostimulation increases fentanyl self-administration in male rats, suggesting stronger motivational or reinforcing effects of fentanyl when paired with optogenetic activation of the CeA. If this is the case, then CeA stimulation should bias choice of fentanyl over an alternative in choice settings. To test this hypothesis, rats were allowed to respond for rewards available on either the left or right lever. In each session, after forced choice trials to ensure subjects were aware of the two options, rats could earn either reinforcer. Once a reinforcer was earned, neither reinforcer was available until a 20s timeout had ended. When fentanyl and fentanyl paired with 0.5s CeA optostimulation (7-10mW, 25HZ; fentanyl with laser) were concurrently available, rats almost exclusively chose fentanyl with laser (Figure 4A, Main effect of reward: F_1,11_=136.5, p<0.0001). The effect of sex or interaction between sex and reward did not reach statistical significance (Sex: F_1,11_=3.68, p=0.0814; Sex x Reward: F_1,11_=4.119, p=0.0673).

**Figure 4.**
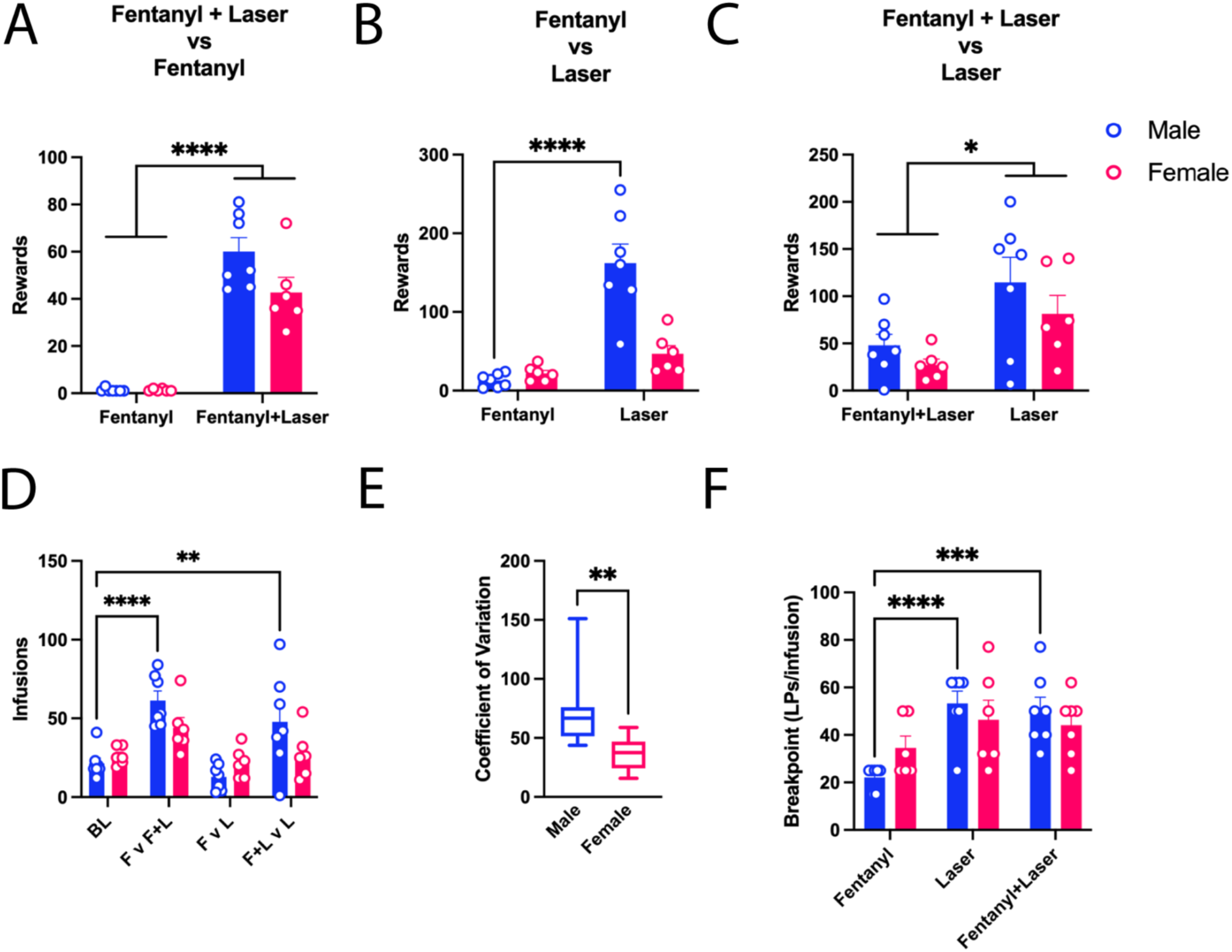
CeA stimulation outcompetes fentanyl as a reinforcer in males (A) Number of Fentanyl versus Fentanyl + Laser reinforcers earned (B) Number of Fentanyl versus Laser reinforcers earned. (C) Number of Fentanyl + Laser versus Laser reinforcers earned. (D)Total fentanyl intake during choice sessions. (D) Coefficient of variation for total fentanyl intake during choice sessions. (F) Progressive ratio breakpoints. Bars represent mean± standard error of the mean. N=6 female, 7 male. * indicates significant effect of reward, sex, or significant pairwise difference, *p<0.05, ** p≤0.01, ***p≤0.001, ****p≤0.0001. Data are presented as mean ± SEM. BL: baseline, F v F+L: Fentanyl vs. Fentanyl + Laser, F v L: Fentanyl vs. Laser, F+L v L: Fentanyl + Laser vs. Laser

It is possible the CeA stimulation is enhancing the motivational or reinforcing effects of the fentanyl. Alternatively, rats might find the CeA stimulation reinforcing in its own right (Fraser et al., 2024). Therefore, we next asked how rats would allocate their behavior between fentanyl on one lever and 0.5s CeA stimulation (7-10mW, 25HZ; laser) on the other. Rats avidly responded for optogenetic stimulation on one lever when fentanyl was available on the other lever, but in a sex-dependent manner (Figure 4B; main effect of reward type (fentanyl vs. optostimulation) and sex on number of rewards earned (Figure 4B, Reward type: F_1,11_=31.52, p=0.0002; Sex: F_1,11_=16.02, p=0.0021; sex x reward type (Sex x Reward type interaction: F_1,11_=16.73, p=0.0018). Males earned significantly more optostimulation rewards than fentanyl rewards (p<0.0001), while females earned similar numbers of each reward type (p=0.504). While there was no significant difference in the number of fentanyl rewards earned between males and females (p=0.900), males earned significantly more optostimulation rewards (p<0.0001). These findings show that males find CeA optostimulation more reinforcing than fentanyl alone at the parameters used here. However, it remains possible that additive effects of CeA optostimulation and fentanyl could be observed, rendering this combination more potent than either optostimulation or fentanyl alone. Therefore, we asked whether Fentanyl with laser would be preferred over laser. In contrast to our expectation, rats persisted in choosing laser (Figure 4C, Reward: F_1,11_=6.954, p=0.0231), in this case, with no sex difference (main effect of Sex: F_1,11_=4.542, p=0.0565; sex x reward type interaction: Reward: F_1,11_=0.0734, p=0.7915).

Interestingly, when we compared the total fentanyl infusions per session, including infusions paired with laser, across the three choice sessions and the baseline session, we found that there was a significant main effect of reward option, i.e. fentanyl as the sole reward compared to choice between fentanyl and laser, and an interaction between reward options and sex (Figure 4D, Reward options: F_3,33_=18.52, p<0.0001; Sex: F_1,11_=0.9178, p=0.3586, Reward options x Sex: F_3,33_=4.297, p=0.0115). Fentanyl intake in males was significantly increased from baseline when the options included fentanyl paired with laser versus laser alone (p=0.0038) or fentanyl paired with laser versus fentanyl alone (p<0.0001). Fentanyl intake was not significantly different from baseline when male rats were allowed to choose between fentanyl and laser (p=0.7899). In females, the difference between baseline intake and fentanyl intake during choice between fentanyl paired with laser and fentanyl alone was not significant (p=0.1371).

Females did not significantly alter fentanyl intake between baseline and fentanyl versus laser (p=0.9937) or between baseline intake and fentanyl paired with laser versus laser alone (p>0.9999). We found that the coefficient of variation of fentanyl intake across baseline and choice sessions was significantly lower in females than males (Figure 4E, Mann-Whitney test: p=0.0047). These findings indicate that optogenetic stimulation of the CeA tends to be more reinforcing than intravenous fentanyl for males, whereas fentanyl intake in females remains more consistent despite the option of direct CeA stimulation.

### CeA stimulation increases motivation for fentanyl in males, but not above that elicited by CeA stimulation alone

To understand how CeA stimulation impacts motivation to respond for fentanyl, rats underwent progressive ratio tests with fentanyl, laser, and fentanyl with laser in which the number of responses required to earn the available reward increased exponentially trial to trial. Analysis revealed a main effect of reward type as well as an interaction between reward type and sex on progressive ratio breakpoint, the maximum number of responses subjects were willing to make in a given session (Figure 4F, Reward: F_1.795,19.74_=14.77, p=0.0002; Reward x Sex: F_2,22_=3.706, p=0.041; Sex: F_1,10_=0.0022, p=0.837). In males, willingness to increase the number of responses for CeA stimulation was higher than fentanyl alone (p=0.0151) and when combined with fentanyl, CeA stimulation significantly increased the number of responses compared to fentanyl (p=0.0072), but there was no difference between breakpoints for fentanyl with laser versus laser alone (p=0.971). In females there was no difference in breakpoints across the three conditions (Fentanyl vs Laser: p=0.166; Fentanyl vs Fentanyl + Laser, p=0.274; Laser vs Fentanyl + Laser, p=0.824), nor did females not significantly differ from males in breakpoints achieved within each outcome (Fentanyl: p=0.164; Fentanyl with laser, p=0.565; Laser, p=0.872). No effect of laser on motivation for fentanyl was observed in eYFP control group (n=5) that did not express ChR2 (supplemental figure S1C).

### Rats persist in seeking optostimulation despite delays even when an alternative reinforcer, fentanyl, is available with no delay

One key difference between IV fentanyl infusion and optogenetic stimulation of CeA neural activity that may account for preference is the timing of the onset of action of the outcomes. While optogenetic activation starts immediately and lasts only 0.5s, fentanyl is infused over 2.8s and its impact on CeA neural activity is constrained by the pharmacokinetics of distribution from blood to brain (Hug Jr. & Murphy, 1981). The onset of action of fentanyl is difficult to accurately determine, but likely occurs in less that 30 seconds (Koyyalagunta, 2007). Still, peak fentanyl reward is delayed when compared to stimulation. To examine whether the onset of action might bias preference towards CeA optostimulation, rats experienced a choice session in which, after sampling trials, a progressive delay was added for delivery of laser rewards such that the first free-choice laser delivery had a 0s delay, the second had a 1s delay, and so on, continuing to increase by one second for every laser reward.

Performance in fentanyl versus laser with progressive delivery delay was compared to the initial fentanyl versus laser (Figure 4B) to determine the effect of delay in laser onset on lever preference across the session. Laser preference decreased as a result of the delay relative to previous fentanyl vs laser performance (Figure 5B, Main effect of delay: F_1,11_=35.77, p<0.001), but the change in fentanyl rewards earned failed to reach statistical significance (Fentanyl rewards earned: No Laser Delay: 17.83±2.788, Laser Delay: 22.75±1.657; Main effect of delay: F_1,11_=4.217, p0.065), indicating rats were not increasing fentanyl intake as they decreased Laser reward. While there was a main effect of sex on laser preference (Sex: F_1,11_=11.78, p=0.0056), the analysis revealed no interaction between sex and delayed laser delivery (Sex X delay: F_1,11_=2.134, p=0.172). The pattern of fentanyl intake was not affected by the progressive delay on the laser reward (Figure 5C, Effect of Delay: F_1,12_=0.0002447, p=0.9878; Effect of Bin: F_23,276_=68.71, p<0.0001; Bin X Delay: F_23,276_=0.4638 p=0.9846). Further, we found no correlation between maximum delay and fentanyl intake (Data not shown, Spearman: r = -0.049, p=0.88). This indicates that there is somewhat independent control of intake of the two reward options, such that a decrease in value of stimulation does not necessarily lead to a compensatory increase in fentanyl value, i.e., rats defend a minimal level of fentanyl intake. Males did appear more likely to increase fentanyl intake, though the interaction between sex and delay condition was not significance (F_1,11_=4.217, p=0.0646). Consistent with our other results, females tended to reach lower maximum delays (Figure 5A, Median Delay: Male: 25s, Female:15s; Effect of Sex: Log-rank (Mantel-Cox) *χ*^2^=4.456, p=0.0348).

**Figure 5.**
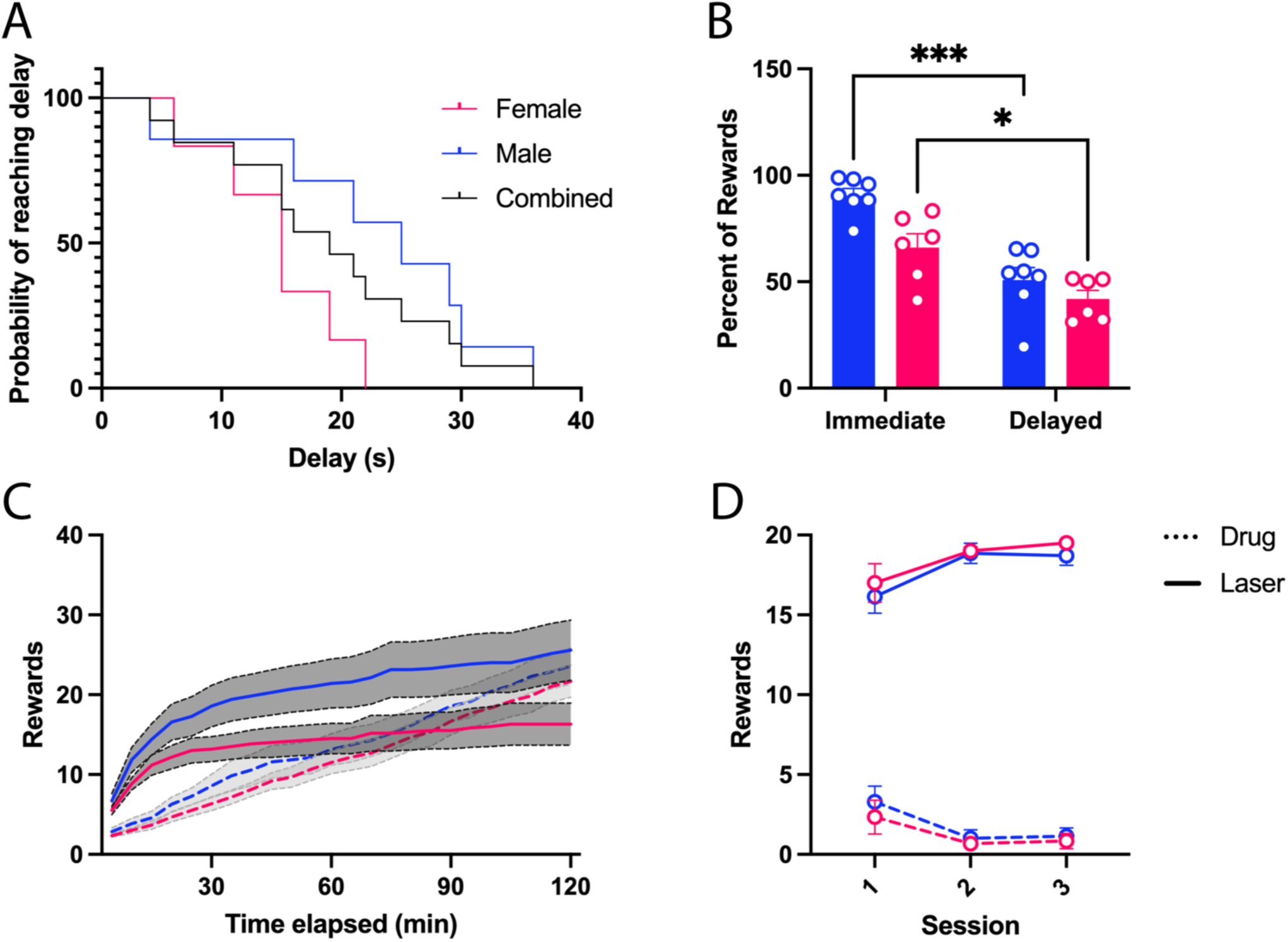
CeA stimulation is pursued despite delays or infrequent rewards (A) Kaplan-meier curve showing probability of rats reaching a given delay in delivery of Laser, Median Delay: Male: 25s, Female:15s (B) Comparison of Laser preference when delivery of Laser is progressively delayed compared to previous choice session with Fentanyl vs. Laser without delay. (C) Cumulative Laser and Fentanyl earned in male and female rats when delivery of laser is progressively delayed, binned in 5-minute increments. Solid lines indicate Laser rewards, dotted lines represent Fentanyl rewards (D) Number of Fentanyl and Laser rewards earned in discrete trial sessions with 20 choice trials. Data are presented as mean ± SEM where applicable. * indicates significant main effect of delay on preference. *p<0.05, ***p≤0.001

### CeA stimulation is reinforcing and preferred over IV fentanyl when rewards are limited and infrequent

One key feature of the choice paradigms used above is that there is no limit on reward acquisition other than both reinforcers being unavailable during the reward delivery and timeout period. Consequently, animals could earn subjectively ideal amounts of both reinforcers without suffering a reduction in the levels of one or the other. The pharmacological effects of fentanyl may also constrain intake while a similar phenomenon of ‘satiety’ does not appear to occur with laser. To address this issue, we used a discrete trial procedure in which rats choose between the fentanyl or laser lever, after which both levers retracted and a 3-min intertrial interval (ITI) ensued until the next trail was initiated by insertion of both levers over three sessions. Given the long ITI between choices, this procedure emphasizes the loss of one possible outcome when the alternative is chosen. The ITI was used to minimize possible impacts of the different durations of action of fentanyl and optogenetic CeA stimulation (0.5s duration).

Rats chose almost exclusively laser rewards (Figure 5D, Main effect of reward: F_1,11_=809.2, p=0.0072). Analysis revealed no main effect of sex or interaction between sex and reward type (Main effect of sex: F_1,11_=0.0639, p=0.805; Sex x reward: F_1,11_=0.926, p=0.357).

### CeA stimulation is highly reinforcing even at low laser powers

The findings above suggest that CeA optogenetic stimulation is itself a potent reinforcer of behavior, potentially obscuring the detection of possible potentiation of fentanyl reward. Thus, we attempted to find a lower power of CeA optostimulation that was no longer reinforcing, but that might still potentiate motivation for a paired reward. We tested the ability of CeA optostimulation at low laser power (0.5-1mW, 0.5s, 25Hz) to support nosepoke responding. Rats nosepoked under FR1 for bilateral optostimulation during 45-minute sessions. No stimulus lights distinguished active from inactive nosepoke port and the active port alternated across sessions to ensure responding was not due to learning in previous sessions.

We observed robust responding (Figure 6A, Stimulations: 845.9±246.5, Active Nosepokes: 1534±569.3, Inactive Nosepokes: 16.7±4.522) that was specific to the active nosepoke (Main effect of nosepoke port: F1,11=8.954, p=0.012) and was independent of sex (Sex: F1,8=1.66, p=0.0.234; Sex x nosepoke port: F1,8=1.69, p=0.230).

**Figure 6.**
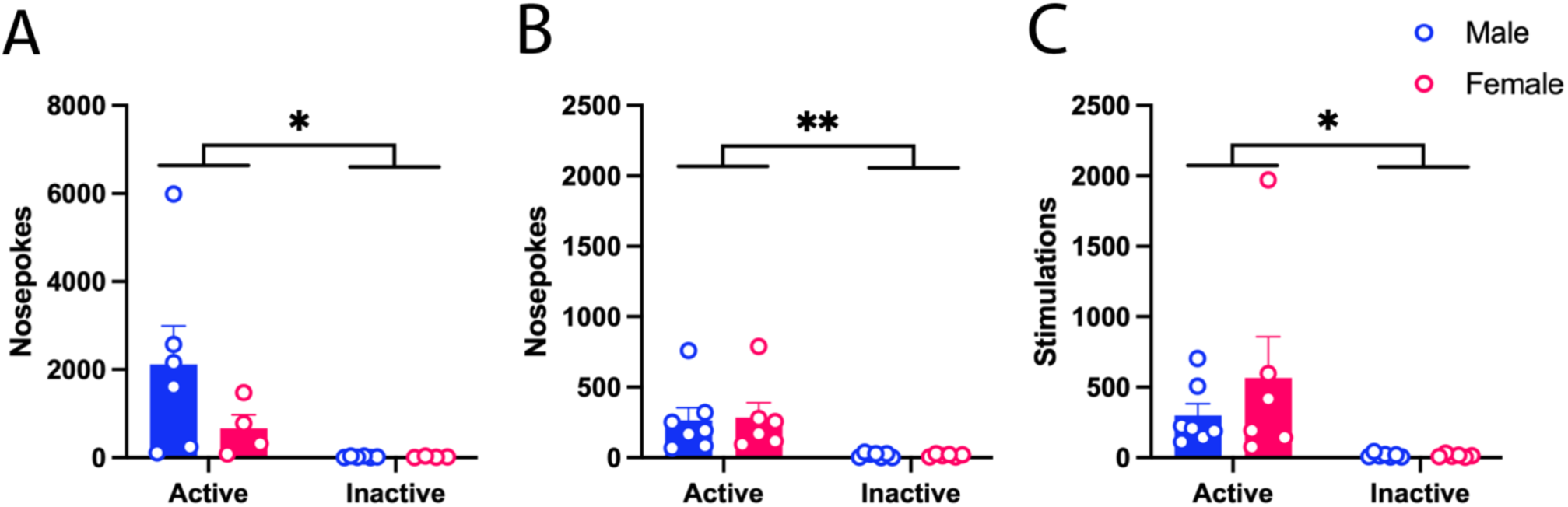
Rats robustly nosepoke for CeA stimulation. Data are presented as mean ± SEM. Active and inactive nosepokes for (A)Bilateral, (B)Left hemisphere, or (C) Right hemisphere CeA stimulation. Bars represent mean± standard error of the mean. Filled magenta symbols are female (n=6), open blue symbols are male (n=6). * indicates significant main effect of nosepoke port, *p<0.05, ** p≤0.01.

Interestingly, the number of optostimulations earned in this task was correlated with the previously determined breakpoint for Fentanyl with laser (Spearman r=0.7347, p=0.018), which indicates that the increase in breakpoint for Fentanyl with laser relates to the potency for optogenetic CeA activation to reinforce behavior. We next repeated these sessions with unilateral stimulation. Rats underwent sessions targeting each hemisphere. Though not as robust, rats reliably acquired responding for unilateral CeA optoactivation despite shifting which port was active (Figure 6B, Left CeA: 186.6±44.07 stimulations, Main effect of nosepoke port; F_1,11_=14.46, p=0.0029; Figure 6C, Right CeA: 286.1±89.03 stimulations, Main effect of nosepoke port: F_1,11_=8.954, p=0.012).

To assess the impact of prior fentanyl self-administration on the reinforcing effects of optogenetic stimulation of the CeA, we assessed the ability of CeA optostimulation to support self-stimulation in fentanyl naïve rats. Fentanyl naïve rats made significantly more entries into the active nosepoke port to earn 500ms blue (473nm) light optogenetic stimulation (25HZ, 0.5-1mW) than the inactive port (Figure S1D, Wilcoxon test: p=0.031). Additionally, when compared to fentanyl experienced rats there was a significant effect of nosepoke port on responding (Active vs. inactive NP: F_(1,10)_=12.38, p=0.006) however there was no significant effect of fentanyl exposure (Main effect of fentanyl exposure: F_(1,10)_=12.38, p=0.897) or interaction between response port and fentanyl experience (Fentanyl exposure x response port:F_(1,10)_=1.038, p=0.332).

Next, fentanyl naïve rats were trained to lever press for optogenetic stimulation of 500ms and then 8s (25HZ, 7-10mW) under an FR3 response requirement. Rats responded significantly more on the active lever when earning 500ms stimulation (Figure S1E, Paired t-test: p=0.025) or 8s stimulation (Not shown, Paired t-test: p=0.007). Further, when fentanyl naïve rats were assessed under a progressive ratio requirement, their breakpoints for optogenetic stimulation did not significantly differ from fentanyl experienced animals (Figure S1F, Mann-Whitney: p=0.266). These findings show potent reinforcing effects of anterior CeA optogenetic stimulation in rats.

## Discussion

Our results demonstrate that direct CeA stimulation is highly reinforcing in both sexes and is consistently preferred over fentanyl in male rats. Previous studies have suggested a critical role for the CeA in the development of the disproportionate motivation for one reward over others, a key component of substance use disorders (DSM-5 Task Force., 2013; Robinson et al., 2014a; Tom et al., 2019a; Warlow et al., 2017, 2020a). Additional studies have demonstrated that the CeA is essential to drug self-administration, craving, resistance to punished drug seeking, and relapse (Barak et al., 2013; Cain et al., 2008c; Chevrette et al., 2002; X. Li et al., 2015b; Marcinkiewcz et al., 2009; Roura-Martínez et al., 2020; W. Sun & Yuill, 2020; Venniro et al., 2020; Wenzel et al., 2011; Xue et al., 2012b; Zhou et al., 2013).

Consistent with previous work assessing the role of the CeA in drug reinforcement, we demonstrate that pharmacological inactivation of the CeA reduces fentanyl intake. We further show that, while pairing CeA stimulation with fentanyl increases fentanyl self-administration and motivation for the reward, motivation for fentanyl with CeA stimulation is not significantly different from motivation for CeA stimulation alone. Further, we demonstrate that CeA stimulation is pursued despite mounting effort requirements, delayed or infrequent delivery, and the presence of a more readily accessible intravenous fentanyl reward. Females were more likely to defend their intake of fentanyl. Though CeA stimulation supported intracranial self-stimulation in both sexes, optostimulation had a decreased effect on fentanyl self-administration and was less preferred over fentanyl in female rats, suggesting that females were more motivated for fentanyl than males. These findings support a multifaceted role for the CeA in substance use disorders, consistent with a larger role in conditioned and unconditioned positively-motivated behaviors (Calu et al., 2010; Fraser et al., 2024; Gallagher et al., 2003; Haney et al., 2010; Holland et al., 2001; Holland & Gallagher, 1993, 1999; Knight et al., 2020; Ross et al., 2016a; Shabel & Janak, 2009; Steinberg et al., 2020a).

### Transferred value of CeA stimulation alters responding for fentanyl

Consistent with previous studies (Robinson et al., 2014b; Warlow et al., 2017, 2020b), CeA stimulation at the time of fentanyl infusion appeared to amplify responding for fentanyl when it was the only reinforcer, biasing responding almost exclusively towards one fentanyl reward over another. Surprisingly, rats were equally motivated to work for CeA stimulation as for the combination of CeA stimulation and fentanyl, suggesting that increased motivation and responding for fentanyl with paired CeA stimulation was determined by the value of CeA stimulation alone rather than a potentiation of the value of fentanyl.

We also observed that only male rats pursued CeA stimulation over fentanyl under free response conditions. Rats had sufficient time to acquire their typical baseline levels of fentanyl in addition to earning CeA stimulations and remained far from any dangerous or excessively sedating fentanyl consumption, yet male rats consistently consumed less fentanyl when CeA stimulation was available. Notably, male rats spent a disproportionate amount of time earning CeA stimulation, despite delays, fentanyl availability, or infrequent availability (see below for discussion on sex differences).

There are many factors that differed between the two options here that could sway choice such as delivery speed, reward magnitude, duration of action. The onset of action of IV fentanyl lags slightly relative to CeA stimulation and it, presumably, has a longer duration of action, including its sedative effects, which limits total intake somewhat (Comer & Cahill, 2018; Koyyalagunta, 2007). These features may contribute to preference for CeA stimulation; however, rats robustly increase fentanyl intake when accompanied by CeA stimulation, indicating that rats are not reaching an intake ceiling in choice sessions. Critically, rats are willing to experience relatively long delays for CeA stimulation, indicating the time-course of the fentanyl effect was not a key factor in biasing subjects towards choosing stimulation.

Despite having many sessions of fentanyl self-administration, sessions were 2 hours long, which is not typically associated with dependence. While there is some evidence that dependence increases motivation for opioids, progressive ratio break points for IV fentanyl in our animals were consistent with those observed in dependent animals in previous studies using long-access self-administration models (6-23hrs), indicating that rats in our experiments did not have low fentanyl motivation due to short access(Awasaki et al., 1997; Fredriksson et al., 2020; Rocío et al., 1999; Stafford et al., 1998; Vasquez et al., 2021; Wade et al., 2015; Yanagita, 1973). The CeA has also been implicated in negative reinforcement of opioids in dependent animals (Chaudun et al., 2024), thus future studies comparing these behaviors in dependent and non-dependent animals are warranted to determine if the role of the CeA might shift.

### Sex Differences

While male and female rats did not differ in the effects of pharmacological manipulations of the CeA or in their CeA self-stimulation behavior, we found sex differences in several tests when stimulation was either paired with fentanyl or pitted against fentanyl. In general, optostimulation had less effect on fentanyl consumption and motivation for fentanyl in females. In fentanyl versus optostimulation choice, female rats earned far fewer stimulations and consequently had less preference for optostimulation compared with males. Similarly, they had lower tolerance for delayed optostimulation when fentanyl was available. Past literature has demonstrated lower demand elasticity in females for fentanyl and cocaine (Andrew Townsend et al., 2019; Kawa & Robinson, 2019), as well as greater drug preference in drug versus non-drug choice (Andrew Townsend et al., 2019; Kerstetter et al., 2012; Lynch, 2018). Though not significant, potentially due to sample size, females in our study appeared to consume greater amounts of fentanyl and achieve higher breakpoints in progressive ratio sessions for fentanyl. This is also consistent with literature showing higher intake and earlier acquisition of self-administration of heroin and cocaine in female rats (Lynch, 2018; Lynch & Carroll, 1999). In line with work demonstrating lower fentanyl demand elasticity in females (Andrew Townsend et al., 2019), females may develop more rigid self-administration behavior and higher motivation for drug, thus optostimulation has less impact on fentanyl self-administration. Females in our study demonstrated significantly lower coefficients of variation in fentanyl intake after acquisition as well as during choice sessions, indicating that females did maintain more rigid fentanyl self-administration. Of note, when opportunities for fentanyl were infrequent, and so a desired set-point of fentanyl intake was unattainable, female rats showed strong preference for CeA optostimulation indistinguishable from males.

It may be that pre-existing or fentanyl dependent sexual dimorphism within the CeA accounts for the effects observed here. Given that females showed similar motivation for CeA stimulation, it does not appear that the sex differences we observed can be explained by differences in reinforcing effects of optostimulation. The sex differences we observed could be due to differences within the CeA and/or in other brain regions, but we do note that prior work found greater sensitivity of CeA neurons in males to alcohol (Hitzemann et al., 2022; Logrip et al., 2017). Of note, corticotropin releasing factor receptor 1 neurons in the CeA also show sex differences in sensitivity to acute ethanol and in response to corticotropin releasing factor, despite comparable baseline properties (Agoglia et al., 2020). Future work further exploring sex differences in CeA involvement in reward and motivation for drugs and natural reward will be important in developing a more translationally-accurate understanding of the CeA in addiction.

We recently reported robust reinforcing properties of CeA optogenetic stimulation in males (Fraser et al., 2024) and here extend that finding to females. Our virus and optical fiber placements in this study were evenly distributed across the three CeA subdivisions (CeC, CeM, and CeL) however they were primarily located within the anterior half of the CeA. The CeA differs along the anterior-posterior (AP) axis both in expression of various markers and previous studies have demonstrated that the effect of some manipulations varies depending on AP axis as well (Bowen et al., 2020; Kong & Zweifel, 2021; Mahler & Berridge, 2009). It is possible that the anatomical subregions of the CeA can account for the failure to obtain robust responding for CeA optogenetic activation in rats in some prior studies (Agüera et al., 2016; Baumgartner et al., 2021, 2022; Knapska et al., 2006; Robinson et al., 2014a; Ross et al., 2016b; Steinberg et al., 2020b; Touzani & Velley, 1998; Valdez et al., 2002; Warlow et al., 2017). Warlow et al. (2020) show that, while unreliable, CeA stimulation does support ICSS in some rats and, in those rats, placement of fibers appears to be restricted to the anterior CeA (Warlow et al., 2020a). Further, data demonstrating the role of the CeA in punished cocaine seeking appears to target primarily the posterior CeA, which may indicate that positive and negative valence are differentially represented along the AP axis (W. L. Sun & Yuill, 2020; Xue et al., 2012c). In mice, robust responding for optogenetic stimulation is observed in a cell type- and projection-specific manner (Kim et al., 2017; Torruella-Suárez et al., 2020). Cell types and projections that support self-stimulation, including projections to the parabrachial nucleus (PBN) and substantia nigra (SN) are enriched in the anterior and intermediate CeA (Baumgartner et al., 2021, 2022; Douglass et al., 2017; Hardaway et al., 2019; Kong & Zweifel, 2021; McCullough et al., 2018; Steinberg et al., 2020b; Torruella-Suárez et al., 2020). Given the complex organization of cell types within the CeA as well as projections to and from the CeA, future work more closely examining distinct CeA subpopulations and projections will be critical in understanding circuit mechanisms within this complex structure.

## Summary

Here we have demonstrated that stimulating neuronal activity within the anterior CeA is so reinforcing that male rats pursue it with intense focus that can lead them to disregard the potent opioid, fentanyl. Male rats consistently preferred CeA stimulation over fentanyl. In contrast, female rats appeared to more strongly defend a minimum fentanyl intake regardless of the alternative rewards available. Our results demonstrate that optogenetic stimulation of neuronal ensembles within the anterior CeA produce a reinforcing effect that inspires compulsive-like responding, but this effect was not transferred to associated rewards. While male and female rats showed similar motivation for CeA stimulation, CeA stimulation as an alternative or added reinforcer altered behavior differently in the two sexes, indicating that there may be important sex differences in the neurobiological basis of opioid use disorders.

## References

1. Agoglia, A. E., Tella, J., & Herman, M. A. (2020). Sex differences in corticotropin releasing factor peptide regulation of inhibitory control and excitability in central amygdala corticotropin releasing factor receptor 1-neurons. Neuropharmacology, 180(April), 108296. 10.1016/j.neuropharm.2020.108296

2. Agüera, A. D. R., García, R., & Puerto, A. (2016). Differential effects of naloxone on rewarding electrical stimulation of the central nucleus of the amygdala and parabrachial complex in a place preference study. Brain Research Bulletin, 124, 182–189. 10.1016/j.brainresbull.2016.04.021

3. Andrew Townsend, E., Stevens Negus, S., Barak Caine, S., & Thomsen, M. (2019). Sex Differences in Opioid Reinforcement Under a Fentanyl vs. Food Choice Procedure in Rats. Neuropsychopharmacology. 10.1038/s41386-019-0356-1

4. Awasaki, Y., Nishida, N., Sasaki, S., & Sato, S. (1997). Dopamine D1 antagonist SCH23390 attenuates self-administration of both cocaine and fentanyl in rats. Environmental Toxicology and Pharmacology, 3(2), 115–122. 10.1016/S1382-6689(97)00147-6

5. Barak, S., Liu, F., Hamida, S. Ben, Yowell, Q. V., Neasta, J., Kharazia, V., Janak, P. H., & Ron, D. (2013). Disruption of alcohol-related memories by mTORC1 inhibition prevents relapse. Nature Neuroscience, 16(8), 1111–1117. 10.1038/nn.3439

6. Baumgartner, H. M., Granillo, M., Schulkin, J., & Berridge, K. C. (2022). Corticotropin releasing factor (CRF) systems: Promoting cocaine pursuit without distress via incentive motivation. PLoS ONE, *17*(5 May), 1–26. 10.1371/journal.pone.0267345

7. Baumgartner, H. M., Schulkin, J., & Berridge, K. C. (2021). Activating Corticotropin-Releasing Factor Systems in the Nucleus Accumbens, Amygdala, and Bed Nucleus of Stria Terminalis: Incentive Motivation or Aversive Motivation? Biological Psychiatry, 89(12), 1162–1175. 10.1016/j.biopsych.2021.01.007

8. Berridge, K. C., & Robinson, T. E. (2016). Liking, Wanting and the Incentive-Sensitization Theory of Addiction. American Psychology, 71(8), 670–679. 10.1037/amp0000059

9. Bowen, A. J., Chen, J. Y., Huang, Y. W., Baertsch, N. A., Park, S., & Palmiter, R. D. (2020). Dissociable control of unconditioned responses and associative fear learning by parabrachial cgrp neurons. ELife, 9, 1–50. 10.7554/ELIFE.59799

10. Cai, Y. Q., Wang, W., Hou, Y. Y., Zhang, Z., Xie, J., & Pan, Z. Z. (2013a). Central amygdala GluA1 facilitates associative learning of opioid reward. Journal of Neuroscience, 33(4), 1577– 1588. 10.1523/JNEUROSCI.1749-12.2013

11. Cai, Y. Q., Wang, W., Hou, Y. Y., Zhang, Z., Xie, J., & Pan, Z. Z. (2013b). Central amygdala GluA1 facilitates associative learning of opioid reward. Journal of Neuroscience, 33(4), 1577– 1588. 10.1523/JNEUROSCI.1749-12.2013

12. Cain, M. E., Denehy, E. D., & Bardo, M. T. (2008a). Individual differences in amphetamine self-administration: The role of the central nucleus of the amygdala. Neuropsychopharmacology, 33(5), 1149–1161. 10.1038/sj.npp.1301478

13. Cain, M. E., Denehy, E. D., & Bardo, M. T. (2008b). Individual differences in amphetamine self-administration: The role of the central nucleus of the amygdala. Neuropsychopharmacology, 33(5), 1149–1161. 10.1038/sj.npp.1301478

14. Cain, M. E., Denehy, E. D., & Bardo, M. T. (2008c). Individual differences in amphetamine self-administration: The role of the central nucleus of the amygdala. Neuropsychopharmacology, 33(5), 1149–1161. 10.1038/sj.npp.1301478

15. Cain, M. E., Denehy, E. D., & Bardo, M. T. (2008d). Individual differences in amphetamine self-administration: The role of the central nucleus of the amygdala. Neuropsychopharmacology, 33(5), 1149–1161. 10.1038/sj.npp.1301478

16. Calu, D. J., Roesch, M. R., Haney, R. Z., Holland, P. C., & Schoenbaum, G. (2010). Neural correlates of variations in event processing during learning in central nucleus of amygdala. Neuron, 68(5), 991–1001. 10.1016/j.neuron.2010.11.019

17. Chaudun, F., Python, L., Liu, Y., Hiver, A., Cand, J., Kieffer, B. L., Valjent, E., & Lüscher, C. (2024). Distinct µ-opioid ensembles trigger positive and negative fentanyl reinforcement. Nature 2024 630:8015, 630(8015), 141–148. 10.1038/s41586-024-07440-x

18. Chevrette, J., Stellar, J. R., Hesse, G. W., & Markou, A. (2002). Both the shell of the nucleus accumbens and the central nucleus of the amygdala support amphetamine self-administration in rats. Pharmacology Biochemistry and Behavior, 71(3), 501–507. 10.1016/S0091-3057(01)00686-4

19. Comer, S. D., & Cahill, C. M. (2018). Fentanyl: Receptor pharmacology, abuse potential, and implications for treatment. Neuroscience & Biobehavioral Reviews. 10.1016/J.NEUBIOREV.2018.12.005

20. de Guglielmo, G., Crawford, E., Kim, S., Vendruscolo, L. F., Hope, B., Brennan, M., Cole, M., Koob, G. F., & George, O. (2016). Recruitment of a Neuronal Ensemble in the Central Nucleus of the Amygdala Is Required for Alcohol Dependence. 10.1523/JNEUROSCI.1395-16.2016

21. de Guglielmo, G., Kallupi, M., Pomrenze, M. B., Crawford, E., Simpson, S., Schweitzer, P., Koob, G. F., Messing, R. O., & George, O. (2019). Inactivation of a CRF-dependent amygdalofugal pathway reverses addiction-like behaviors in alcohol-dependent rats. Nature Communications, 10(1), 1238. 10.1038/s41467-019-09183-0

22. Domi, E., Xu, L., Toivainen, S., Nordeman, A., Gobbo, F., Venniro, M., Shaham, Y., Messing, R. O., Visser, E., van den Oever, M. C., Holm, L., Barbier, E., Augier, E., & Heilig, M. (2021). A neural substrate of compulsive alcohol use. Science Advances, 7(34). 10.1126/sciadv.abg9045

23. Douglass, A. M., Kucukdereli, H., Ponserre, M., Markovic, M., Gründemann, J., Strobel, C., Alcala Morales, P. L., Conzelmann, K. K., Lüthi, A., & Klein, R. (2017). Central amygdala circuits modulate food consumption through a positive-valence mechanism. Nature Neuroscience, 20(10), 1384–1394. 10.1038/nn.4623

24. DSM-5 Task Force. (2013). Diagnostic and statistical manual of mental disorders : DSM-5. American Psychiatric Association.

25. Fragale, J. E., James, M. H., & Aston-Jones, G. (2020). Intermittent self-administration of fentanyl induces a multifaceted addiction state associated with persistent changes in the orexin system. BioRxiv, 2020.04.23.055848. 10.1101/2020.04.23.055848

26. Fraser, K. M., Kim, T. H., Castro, M., Drieu, C., Padovan-Hernandez, Y., Chen, B., Pat, F., Ottenheimer, D. J., & Janak, P. H. (2024). Encoding and context-dependent control of reward consumption within the central nucleus of the amygdala. IScience, 27(5), 109652. 10.1016/j.isci.2024.109652

27. Fredriksson, I., Applebey, S. V., Minier-Toribio, A., Shekara, A., Bossert, J. M., & Shaham, Y. (2020). Effect of the dopamine stabilizer (-)-OSU6162 on potentiated incubation of opioid craving after electric barrier-induced voluntary abstinence. Neuropsychopharmacology, 45(5), 770–779. 10.1038/s41386-020-0602-6

28. Gallagher, M., Graham, P. W., & Holland, P. C. (2003). The Amygdala Central Nucleus and Appetitive Pavlovian Conditioning: Lesions Impair One Class of Conditioned Behavior. J. Neurosci, 10(6), 1–6.

29. Haney, R. Z., Calu, D. J., Takahashi, Y. K., Hughes, B. W., & Schoenbaum, G. (2010). Inactivation of the central but not the basolateral nucleus of the amygdala disrupts learning in response to overexpectation of reward. The Journal of Neuroscience : The Official Journal of the Society for Neuroscience, 30(8), 2911–2917. 10.1523/JNEUROSCI.0054-10.2010

30. Hardaway, J. A., Halladay, L. R., Mazzone, C. M., Pati, D., Bloodgood, D. W., Kim, M., Jensen, J., DiBerto, J. F., Boyt, K. M., Shiddapur, A., Erfani, A., Hon, O. J., Neira, S., Stanhope, C. M., Sugam, J. A., Saddoris, M. P., Tipton, G., McElligott, Z., Jhou, T. C., … Kash, T. L. (2019). Central Amygdala Prepronociceptin-Expressing Neurons Mediate Palatable Food Consumption and Reward. Neuron, 102(5), 1037–1052.e7. 10.1016/j.neuron.2019.03.037

31. Hitzemann, R., Bergeson, S. E., Berman, A. E., Bubier, J. A., Chesler, E. J., Finn, D. A., Hein, M., Hoffman, P., Holmes, A., Kisby, B. R., Lockwood, D., Lodowski, K. H., McManus, M., Owen, J. A., Ozburn, A. R., Panthagani, P., Ponomarev, I., Saba, L., Tabakoff, B., … Phillips, T. J. (2022). Sex Differences in the Brain Transcriptome Related to Alcohol Effects and Alcohol Use Disorder. Biological Psychiatry, 91(1), 43–52. 10.1016/j.biopsych.2021.04.016

32. Holland, P. C., Chik, Y., & Zhang, Q. (2001). Inhibitory learning tests of conditioned stimulus associability in rats with lesions of the amygdala central nucleus. Behavioral Neuroscience, 115(5), 1154–1158. 10.1037/0735-7044.115.5.1154

33. Holland, P. C., & Gallagher, M. (1993). Amygdala Central Nucleus Lesions Disrupt Increments, But Not Decrements, in Conditioned Stimulus Processing. Behavioral Neuroscience, 107(2), 246–253. 10.1037/0735-7044.107.2.246

34. Holland, P. C., & Gallagher, M. (1999). Amygdala circuitry in attentional and representational processes. Trends in Cognitive Sciences, 3(2), 65–73. 10.1016/S1364-6613(98)01271-6

35. Hou, Y.-Y., Cai, Y.-Q., & Pan, Z. Z. (2017). GluA1 in central amygdala promotes opioid use and reverses inhibitory effect of pain. Physiology & Behavior, 176(1), 100–106. 10.1177/0022146515594631.Marriage

36. Hug Jr., C. C., & Murphy, M. R. (1981). Tissue Redistribution of Fentanyl and Termination of Its Effects in Rats. Anesthesiology, 55(4), 369–375. 10.1097/00000542-198110000-00006

37. Kallupi, M., Carrette, L. L. G., Kononoff, J., Woods, L. C. S., Palmer, A. A., Schweitzer, P., George, O., & De Guglielmo, G. (2020a). Nociceptin attenuates the escalation of oxycodone self-administration by normalizing CeA-GABA transmission in highly addicted rats. Proceedings of the National Academy of Sciences of the United States of America, 117(4), 2140– 2148. 10.1073/pnas.1915143117

38. Kallupi, M., Carrette, L. L. G., Kononoff, J., Woods, L. C. S., Palmer, A. A., Schweitzer, P., George, O., & De Guglielmo, G. (2020b). Nociceptin attenuates the escalation of oxycodone self-administration by normalizing CeA-GABA transmission in highly addicted rats. Proceedings of the National Academy of Sciences of the United States of America, 117(4), 2140– 2148. 10.1073/pnas.1915143117

39. Kawa, A. B., & Robinson, T. E. (2019). Sex differences in incentive-sensitization produced by intermittent access cocaine self-administration. Psychopharmacology, 236(2), 625–639. 10.1007/s00213-018-5091-5

40. Kerstetter, K. A., Ballis, M. A., Duffin-Lutgen, S., Carr, A. E., Behrens, A. M., & Kippin, T. E. (2012). Sex differences in selecting between food and cocaine reinforcement are mediated by estrogen. Neuropsychopharmacology, 37(12), 2605–2614. 10.1038/npp.2012.99

41. Kim, J., Zhang, X., Muralidhar, S., LeBlanc, S. A., & Tonegawa, S. (2017). Basolateral to Central Amygdala Neural Circuits for Appetitive Behaviors. Neuron, 93(6), 1464–1479.e5. 10.1016/j.neuron.2017.02.034

42. Knapska, E., Walasek, G., Nikolaev, E., Neuhäusser-Wespy, F., Lipp, H.-P., Kaczmarek, L., & Werka, T. (2006). Differential involvement of the central amygdala in appetitive versus aversive learning. Learning & Memory (Cold Spring Harbor, N.Y.), 13(2), 192–200. 10.1101/lm.54706

43. Knight, C. P., Hauser, S. R., Waeiss, R. A., Molosh, A. I., Johnson, P. L., Truitt, W. A., McBride, W. J., Bell, R. L., Shekhar, A., & Rodd, Z. A. (2020). The Rewarding and Anxiolytic Properties of Ethanol within the Central Nucleus of the Amygdala: Mediated by Genetic Background and Nociceptin. Journal of Pharmacology and Experimental Therapeutics, 374(3), 366–375. 10.1124/JPET.119.262097

44. Kong, M. S., & Zweifel, L. S. (2021). Central amygdala circuits in valence and salience processing. Behavioural Brain Research, 410(February), 113355. 10.1016/j.bbr.2021.113355

45. Koob, G. F., & Volkow, N. D. (2010). Neurocircuitry of addiction. Neuropsychopharmacology : Official Publication of the American College of Neuropsychopharmacology, 35(1), 217–238. 10.1038/npp.2009.110

46. Koyyalagunta, D. (2007). Opioid Analgesics. Pain Management, 2, 939–964. 10.1016/B978-0-7216-0334-6.50117-5

47. Li, X., Zeric, T., Kambhampati, S., Bossert, J. M., & Shaham, Y. (2015a). The Central Amygdala Nucleus is Critical for Incubation of Methamphetamine Craving. Neuropsychopharmacology, 40(5), 1297–1306. 10.1038/npp.2014.320

48. Li, X., Zeric, T., Kambhampati, S., Bossert, J. M., & Shaham, Y. (2015b). The Central Amygdala Nucleus is Critical for Incubation of Methamphetamine Craving. Neuropsychopharmacology, 40(5), 1297–1306. 10.1038/npp.2014.320

49. Li, Y. Q., Li, F. Q., Wang, X. Y., Wu, P., Zhao, M., Xu, C. M., Shaham, Y., & Lu, L. (2008). Central amygdala extracellular signal-regulated kinase signaling pathway is critical to incubation of opiate craving. The Journal of Neuroscience : The Official Journal of the Society for Neuroscience, 28(49), 13248–13257. 10.1523/JNEUROSCI.3027-08.2008

50. Logrip, M. L., Oleata, C., & Roberto, M. (2017). Sex differences in responses of the basolateral-central amygdala circuit to alcohol, corticosterone and their interaction. Neuropharmacology, 114, 123–134. 10.1016/j.neuropharm.2016.11.021

51. Lynch, W. J. (2018). Modeling the development of drug addiction in male and female animals. Pharmacology Biochemistry and Behavior, 164, 50–61. 10.1016/J.PBB.2017.06.006

52. Lynch, W. J., & Carroll, M. E. (1999). Sex differences in the acquisition of intravenously self-administered cocaine and heroin in rats. Psychopharmacology, 144(1), 77–82. 10.1007/s002130050979

53. Mahler, S. V., & Berridge, K. C. (2009a). Which cue to “Want?” central amygdala opioid activation enhances and focuses incentive salience on a prepotent reward cue. Journal of Neuroscience, 29(20), 6500–6513. 10.1523/JNEUROSCI.3875-08.2009

54. Mahler, S. V., & Berridge, K. C. (2009b). Which cue to “Want?” central amygdala opioid activation enhances and focuses incentive salience on a prepotent reward cue. Journal of Neuroscience, 29(20), 6500–6513. 10.1523/JNEUROSCI.3875-08.2009

55. Marcinkiewcz, C. A., Prado, M. M., Isaac, S. K., Marshall, A., Rylkova, D., & Bruijnzeel, A. W. (2009). Corticotropin-releasing factor within the central nucleus of the amygdala and the nucleus accumbens shell mediates the negative affective state of nicotine withdrawal in rats. 34(7), 1743–1752. 10.1038/npp.2008.231.Corticotropin-releasing

56. McCullough, K. M., Morrison, F. G., Hartmann, J., Carlezon, W. A., & Ressler, K. J. (2018). Quantified coexpression analysis of central amygdala subpopulations. ENeuro, 5(1). 10.1523/ENEURO.0010-18.2018

57. Paxinos, G., & Watson, C. (2009). The Rat Brain in Stereotaxic Coordinates. Elsevier/Academic.

58. Pelloux, Y., Minier-Toribio, A., Hoots, J. K., Bossert, J. M., & Shaham, Y. (2018). Opposite effects of basolateral amygdala inactivation on context-induced relapse to cocaine seeking after extinction versus punishment. Journal of Neuroscience, 38(1), 51–59. 10.1523/JNEUROSCI.2521-17.2017

59. Robinson, M. J. F., Warlow, S. M., & Berridge, K. C. (2014a). Optogenetic excitation of central amygdala amplifies and narrows incentive motivation to pursue one reward above another. The Journal of Neuroscience : The Official Journal of the Society for Neuroscience, 34(50), 16567– 16580. 10.1523/JNEUROSCI.2013-14.2014

60. Robinson, M. J. F., Warlow, S. M., & Berridge, K. C. (2014b). Optogenetic excitation of central amygdala amplifies and narrows incentive motivation to pursue one reward above another. The Journal of Neuroscience : The Official Journal of the Society for Neuroscience, 34(50), 16567– 16580. 10.1523/JNEUROSCI.2013-14.2014

61. Rocío, M., Carrera, M. R. A., Schulteis, G., & Koob, G. F. (1999). Heroin self-administration in dependent Wistar rats: Increased sensitivity to naloxone. Psychopharmacology, 144(2), 111–120. 10.1007/s002130050983

62. Ross, S. E., Lehmann Levin, E., Itoga, C. A., Schoen, C. B., Selmane, R., & Aldridge, J. W. (2016a). Deep brain stimulation in the central nucleus of the amygdala decreases ‘wanting’ and ‘liking’ of food rewards. European Journal of Neuroscience, 44(7), 2431–2445. 10.1111/ejn.13342

63. Ross, S. E., Lehmann Levin, E., Itoga, C. A., Schoen, C. B., Selmane, R., & Aldridge, J. W. (2016b). Deep brain stimulation in the central nucleus of the amygdala decreases ‘wanting’ and ‘liking’ of food rewards. European Journal of Neuroscience, 44(7), 2431–2445. 10.1111/ejn.13342

64. Roura-Martínez, D., Ucha, M., Orihuel, J., Ballesteros-Yáñez, I., Castillo, C. A., Marcos, A., Ambrosio, E., & Higuera-Matas, A. (2020). Central nucleus of the amygdala as a common substrate of the incubation of drug and natural reinforcer seeking. Addiction Biology, 25(2). 10.1111/adb.12706

65. Shabel, S. J., & Janak, P. H. (2009). Substantial similarity in amygdala neuronal activity during conditioned appetitive and aversive emotional arousal. Proceedings of the National Academy of Sciences, 106(35), 15031–15036. 10.1073/pnas.0905580106

66. Stafford, D., LeSage, M. G., & Glowa, J. R. (1998). Progressive-ratio schedules of drug delivery in the analysis of drug self-administration: A review. Psychopharmacology, 139(3), 169–184. 10.1007/s002130050702

67. Steinberg, E. E., Gore, F., Heifets, B. D., Taylor, M. D., Norville, Z. C., Beier, K. T., Földy, C., Lerner, T. N., Luo, L., Deisseroth, K., & Malenka, R. C. (2020a). Amygdala-Midbrain Connections Modulate Appetitive and Aversive Learning. Neuron, 106(6), 1026–1043.e9. 10.1016/j.neuron.2020.03.016

68. Steinberg, E. E., Gore, F., Heifets, B. D., Taylor, M. D., Norville, Z. C., Beier, K. T., Földy, C., Lerner, T. N., Luo, L., Deisseroth, K., & Malenka, R. C. (2020b). Amygdala-Midbrain Connections Modulate Appetitive and Aversive Learning. Neuron, 106(6), 1026–1043.e9. 10.1016/j.neuron.2020.03.016

69. Sun, W. L., & Yuill, M. B. (2020). Role of the GABAa and GABAb receptors of the central nucleus of the amygdala in compulsive cocaine-seeking behavior in male rats. Psychopharmacology, 237(12), 3759–3771. 10.1007/s00213-020-05653-2

70. Sun, W., & Yuill, M. B. (2020a). Role of the GABA a and GABA b receptors of the central nucleus of the amygdala in compulsive cocaine-seeking behavior in male rats. Psychopharmacology, 237(12), 3759–3771. 10.1007/s00213-020-05653-2/Published

71. Sun, W., & Yuill, M. B. (2020b). Role of the GABA a and GABA b receptors of the central nucleus of the amygdala in compulsive cocaine-seeking behavior in male rats. Psychopharmacology, 237(12), 3759–3771. 10.1007/s00213-020-05653-2/Published

72. Thomsen, M., & Caine, S. B. (2005). Chronic intravenous drug self-administration in rats and mice. In Current protocols in neuroscience / editorial board, Jacqueline N. Crawley … [et al.]: Vol. Chapter 9. 10.1002/0471142301.ns0920s32

73. Tom, R. L., Ahuja, A., Maniates, H., Freeland, C. M., & Robinson, M. J. F. (2019a). Optogenetic activation of the central amygdala generates addiction-like preference for reward. European Journal of Neuroscience, 50(3), 2086–2100. 10.1111/ejn.13967

74. Tom, R. L., Ahuja, A., Maniates, H., Freeland, C. M., & Robinson, M. J. F. (2019b). Optogenetic activation of the central amygdala generates addiction-like preference for reward. European Journal of Neuroscience, 50(3), 2086–2100. 10.1111/ejn.13967

75. Torruella-Suárez, M. L., Vandenberg, J. R., Cogan, E. S., Tipton, G. J., Teklezghi, A., Dange, K., Patel, G. K., McHenry, J. A., Andrew Hardaway, J., Kantak, P. A., Crowley, N. A., DiBerto, J. F., Faccidomo, S. P., Hodge, C. W., Stuber, G. D., & McElligott, Z. A. (2020). Manipulations of central amygdala neurotensin neurons alter the consumption of ethanol and sweet fluids in mice. Journal of Neuroscience, 40(3), 632–647. 10.1523/JNEUROSCI.1466-19.2019

76. Touzani, K., & Velley, L. (1998). Electrical self-stimulation in the central amygdaloid nucleus after ibotenic acid lesion of the lateral hypothalamus. Behavioural Brain Research, 90(2), 115–124. 10.1016/s0166-4328(97)00090-9

77. Valdez, G. R., Roberts, A. J., Chan, K., Davis, H., Brennan, M., Zorrilla, E. P., & Koob, G. F. (2002). Increased Ethanol Self-Administration and Anxiety-Like Behavior During Acute Ethanol Withdrawal and Protracted Abstinence: Regulation by Corticotropin-Releasing Factor. Alcoholism: Clinical and Experimental Research, 26(10), 1494–1501. 10.1111/j.1530-0277.2002.tb02448.x

78. Vasquez, T. E. S., Shah, P., Di Re, J., Laezza, F., & Green, T. A. (2021). Individual differences in frustrative nonreward behavior for sucrose in rats predict motivation for fentanyl under progressive ratio. ENeuro, 8(5), 1–8. 10.1523/ENEURO.0136-21.2021

79. Venniro, M., Russell, T. I., Ramsey, L. A., Richie, C. T., Lesscher, H. M. B., Giovanetti, S. M., Messing, R. O., & Shaham, Y. (2020). Abstinence-dependent dissociable central amygdala microcircuits control drug craving. 117(14), 8126–8134. 10.1073/pnas.2001615117/-/DCSupplemental

80. Wade, C. L., Vendruscolo, L. F., Schlosburg, J. E., Hernandez, D. O., & Koob, G. F. (2015). Compulsive-like responding for opioid analgesics in rats with extended access. Neuropsychopharmacology, 40(2), 421–428. 10.1038/npp.2014.188

81. Warlow, S. M., Naffziger, E. E., & Berridge, K. C. (2020a). The central amygdala recruits mesocorticolimbic circuitry for pursuit of reward or pain. Nature Communications, 11(1), 1–15. 10.1038/s41467-020-16407-1

82. Warlow, S. M., Naffziger, E. E., & Berridge, K. C. (2020b). The central amygdala recruits mesocorticolimbic circuitry for pursuit of reward or pain. Nature Communications, 11(1), 1–15. 10.1038/s41467-020-16407-1

83. Warlow, S. M., Robinson, M. J. F., & Berridge, K. C. (2017). Optogenetic Central Amygdala Stimulation Intensifies and Narrows Motivation for Cocaine. The Journal of Neuroscience : The Official Journal of the Society for Neuroscience, 37(35), 8330–8348. 10.1523/JNEUROSCI.3141-16.2017

84. Wenzel, J. M., Waldroup, S. A., Haber, Z. M., Su, Z. I., Ben-Shahar, O., & Ettenberg, A. (2011). Effects of lidocaine-induced inactivation of the bed nucleus of the stria terminalis, the central or the basolateral nucleus of the amygdala on the opponent-process actions of self-administered cocaine in rats. Psychopharmacology, 217(2), 221–230. 10.1007/s00213-011-2267-7

85. Xue, Y., Steketee, J. D., & Sun, W. (2012a). Inactivation of the central nucleus of the amygdala reduces the effect of punishment on cocaine self-administration in rats. European Journal of Neuroscience, 35(5), 775–783. 10.1111/j.1460-9568.2012.08000.x

86. Xue, Y., Steketee, J. D., & Sun, W. (2012b). Inactivation of the central nucleus of the amygdala reduces the effect of punishment on cocaine self-administration in rats. European Journal of Neuroscience, 35(5), 775–783. 10.1111/j.1460-9568.2012.08000.x

87. Xue, Y., Steketee, J. D., & Sun, W. (2012c). Inactivation of the central nucleus of the amygdala reduces the effect of punishment on cocaine self-administration in rats. European Journal of Neuroscience, 35(5), 775–783. 10.1111/j.1460-9568.2012.08000.x

88. Yanagita, T. (1973). An experimental framework for evaluation of dependence liability of various types of drugs in monkeys. Bulletin on Narcotics, 25, 57–64.

89. Zhou, Y., Leri, F., Grella, S. L., Aldrich, J. V., & Kreek, M. J. (2013). Involvement of dynorphin and kappa opioid receptor in yohimbine-induced reinstatement of heroin seeking in rats. Synapse, 67(6), 358–361. 10.1002/syn.21638

