## Supplemental figure 1 for "Optogenetic Central Amygdala Stimulation is Highly Reinforcing and Strongly Outcompetes Fentanyl Self-Administration in Male, but not Female, Rats"

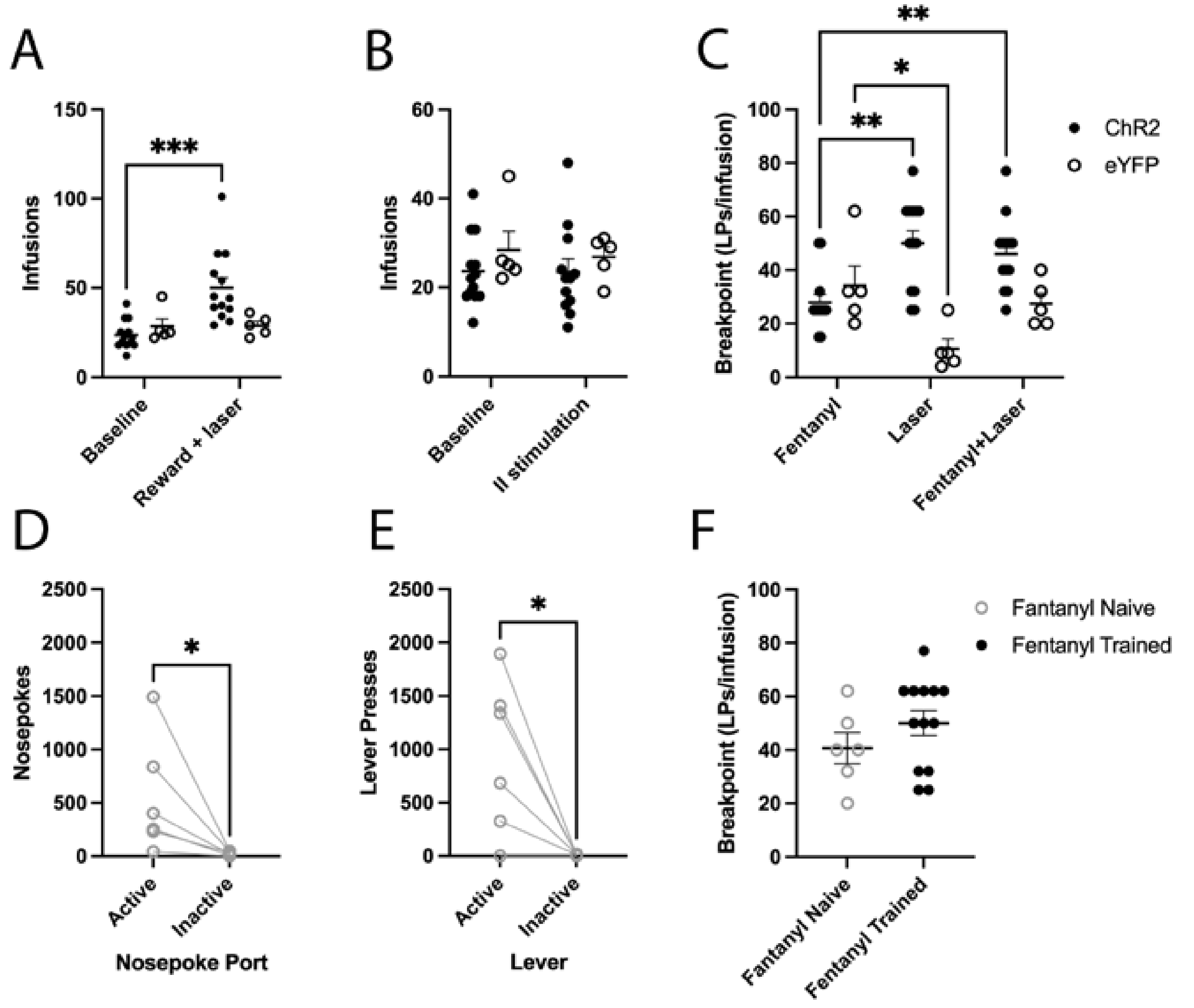


**Figure S1** Optogenetic stimulation of the CeA is reinforcing independent in fentanyl naïve rats Data are presented as mean ± SEM. (A), (B) Comparison of baseline fentanyl intake versus fentanyl intake in a session (A) pairing laser and fentanyl infusion and (B) with laser delivery during inter-infusion intervals in rats with eYFP infused into the CeA (C) Comparison of progressive ratio breakpoints in rats with eYFP infused into the CeA versus ChR2, N=12 ChR2, 5 eYFP. (D) Active and inactive nosepokes for optogenetic CeA stimulation in fentanyl Naïve rats (E) Active and inactive lever presses for optogenetic CeA stimulation in fentanyl Naïve rats under an FR3 response requirement. (F) Comparison of progressive ratio breakpoints in fentanyl naïve rats (n=6) versus fentanyl trained rats (n=12) * indicates significant difference between conditions, *p<0.05, ** p≤0.01.
